# De novo design of potent inhibitors of Clostridioides difficile toxin B

**DOI:** 10.1101/2024.08.26.609740

**Authors:** Robert J. Ragotte, John Tam, Sean Miletic, Roger Palou, Connor Weidle, Zhijie Li, Matthias Glögl, Greg L. Beilhartz, Huazhu Liang, Kenneth D. Carr, Andrew J. Borst, Brian Coventry, Xinru Wang, John L. Rubinstein, Mike Tyers, Roman A. Melnyk, David Baker

## Abstract

*Clostridioides difficile* is a major cause of secondary disease in hospitals. During infection, *C. difficile* toxin B drives disease pathology. Here we use deep learning and Rosetta-based approaches to de novo design small proteins that block the entry of TcdB into cells. These molecules have binding affinities and neutralization IC50’s in the pM range and are compelling candidates for further clinical development. By directly targeting the toxin rather than the pathogen, these molecules have the advantage of immediate cessation of disease and lower selective pressure for escape compared to conventional antibiotics. As *C. difficile* infects the colon, the protease and pH resistance of the designed proteins opens the door to oral delivery of engineered biologics.

**Significance statement:** *C. difficile* infection (CDI) is a major public health concern with over half a million cases in the United States annually resulting in 30,000 deaths. Current therapies are inadequate and frequently result in cycles of recurrent infection (rCDI). Progress has been made in the development of anti-toxin mAb therapies that can reduce the rate of rCDI, but these remain unaffordable and out of reach for many patients. Using de novo protein design, we developed small protein inhibitors targeting two independent receptor binding sites on the toxin that drives pathology during CDI. These molecules are high affinity, potently neutralizing and stable in simulated intestinal fluid, making them strong candidates for the clinical development of new CDI therapies.

## Introduction

*Clostridioides difficile* toxin B (TcdB) drives the pathology of *C. difficile* infection (CDI), a common and sometimes fatal nosocomial disease of the colon. Currently, vancomycin and fidaxomicin remain the standard-of-care for acute CDI, but perturbations of the microbiota can contribute to disease progression(1), resulting in a high rate of relapse and patients entering cycles of recurrent CDI (rCDI). Bezlotoxumab, a therapeutic monoclonal antibody targeting TcdB, reduces the rate of rCDI by 10%, while a neutralizing mAb against TcdA did not show clinical efficacy, providing a strong biological basis for targeting TcdB neutralization during CDI(2, 3). However, given the high cost, complex route of administration (intravenous) and unknown efficacy during acute CDI, bezlotoxumab is only recommended for those at high risk of rCDI and it remains unclear how it compares to fidaxomicin, a comparably priced but orally administered antibiotic that similarly reduces that rate rCDI(4, 5). There is a need for new anti-TcdB therapeutics that do not require intravenous administration, act directly in the site of infection, are affordable and could be prescribed prophylactically during outbreaks without substantial risk of driving antibiotic resistance.

We reasoned that de novo designed miniprotein inhibitors could help address unmet clinical needs for treating CDI. Miniproteins have advantages over monoclonal antibodies in ease of production, thermostability, and protease stability while retaining high affinity and specificity for their targets(6–9). We set out to design small proteins that precisely target sites on TcdB that engage cell surface receptors, thereby preventing the toxin from entering cells.

### Inhibition of the Frizzled/TFPI binding site

Receptor usage patterns of TcdB differ between variants of the toxin(10). With the exception of TcdB2 which primarily uses the CSPG4 binding site(10), TcdB uses a receptor binding site in the distal RBD that can interact with Frizzled-1/2/7(11) or TFPI(12), as illustrated in Figure 1. A second receptor binding site is located in the toxin core where the toxin can bind to host CSPG4(13), used alongside Frizzled-1/2/7 or independent of Frizzled in the case of TcdB2.

**Figure 1.**
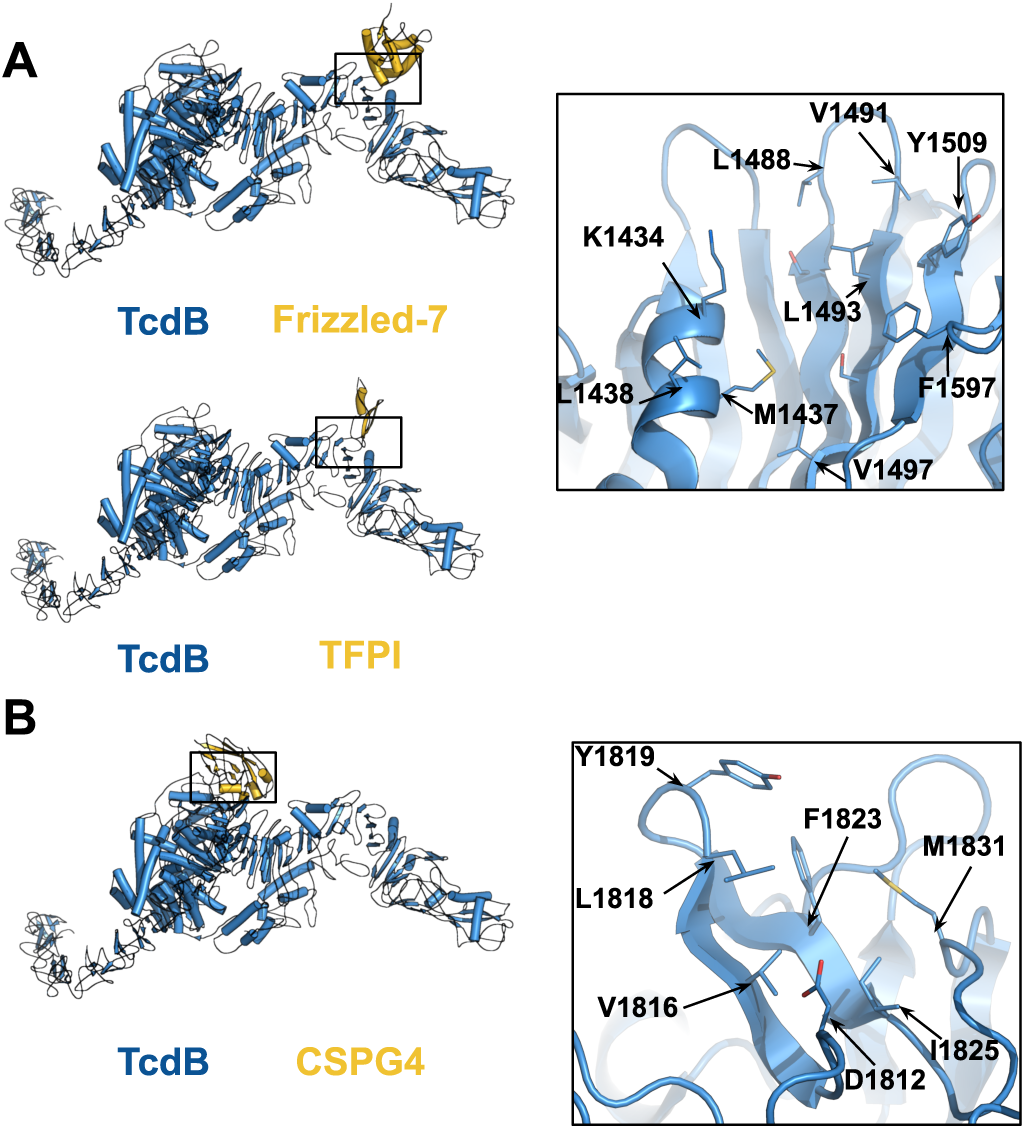
Composite models of TcdB-host cell receptor complexes. Inset boxes highlight the residues targeted during design. A. TcdB interface with Frizzled proteins (1, 2 and 7) and TFPI. B. TcdB interface with CSPG4.

We began by targeting the TcdB Frizzled/TFPI binding interface using a design pipeline integrating Rosetta- and deep learning-based methods(7, 14). We generated docks from 3-helical and 4-helical bundle libraries using RifGen and RifDock focused on hydrophobic residues at the binding interface (Figure 1A). We then assigned sequences to the docks using ProteinMPNN and filtered the designs using AlphaFold2 initial guess(14, 15). 15,000 designs were screened for binding using yeast surface display followed by optimization through site saturation mutagenesis(7) (Figure S1). After sequence optimization, the tightest binders were expressed in *E. coli,* and three sets of designs (each resulting from a single initial dock) had monodisperse SEC traces and high affinity for their target sites (Figure 2A, Figure S2). These are defined as group 1, group 2 and group 3 designs. Designs within each group originate from the same initial dock (Figure 2A).

**Figure 2.**
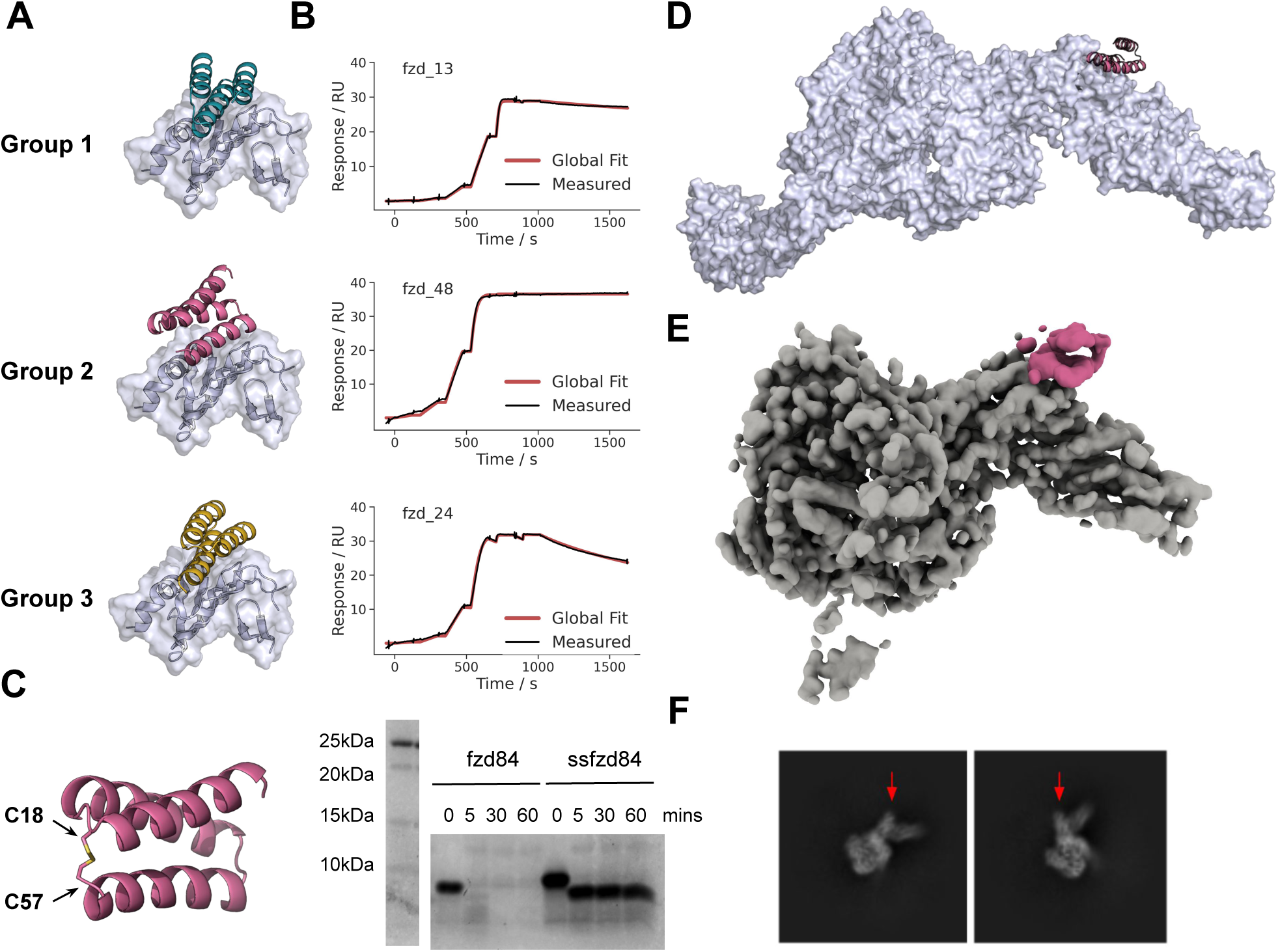
Design of anti-TcdB Frizzled-blocking minibinders. **A.** Design models of three high affinity binder families. **B.** Corresponding SPR traces of a design from each family with TcdB RBD captured on chip and a 6-step 4-fold dilution series starting at 100 nM. **C.** Disulfide stabilization of fzd84 minibinder with the location of the disulfide introduced (left) and a coomassie stained SDS-PAGE after a time course incubation in simulated intestinal fluid (with 0.1 mg/ml of trypsin and chymotrypsin) at 37°C (right). **D**. Design model of fzd48 in complex with the full length toxin from PDB 6OQ5. E. Segmented cryoEM map of TcdB in complex with fzd48 (in pink). F. Example class averages with arrows indicating additional density due to fzd48 binding.

For group 2 designs, accurate Koffs could not be determined even with dissociation times > 1 h, as minimal dissociation was detected (Figure 2B, S5), suggesting K_D_s in the low pM range. In neutralization assays using CSPG4-deficient Vero cells, fzd5 (group 1), fzd48 (group 2) and fzd84 (group 2) had IC50s of 31 (95% CI 22 - 48), 133 (95% CI 97 - 187) and 61 (95% CI 38 - 101) pM in the presence of 1.5 pM toxin respectively (Figure 4A, 4B).

The structure of fzd48 in complex with TcdB was determined by electron cryo-microscopy (cryoEM). The binding epitope and orientation match the design model (Figure 2D-F, S3, Table S4). At 4.6Å resolution, we could clearly resolve the secondary structure of fzd48 with density for all three helices at its intended target site on the knuckle of the RBD (Figure 2D-F). In the design model of fzd48, W47 extends into the hydrophobic pocket formed from the coming together of the kinked α-helix and the 5-stranded β-sheet in the RBD that composes the bulk of the interface. K35 and K54 flank this hydrophobic core of the interface, potentially forming salt bridges with neighboring D1490 and E1547 on TcdB (the limited resolution of the cryoEM map prevented building and assessment of sidechain accuracy).

TcdB acts in the colonic lumen and hence is only amenable for treatment by protein-based biologics if an inhibitor can survive in the gastrointestinal tract without extensive proteolysis. To improve the protease stability of the designs (which were degraded rapidly in simulated intestinal fluid (Figure 2C, S4), we introduced disulfides into the group 1 and group 2 designs. In the group 2 design for example, the disulfide connects a loop between the first and second helix with the C terminus, effectively locking the structure together (Figure 2C). Both group 1 and group 2 disulfide stabilized designs retained their binding and neutralizing activity and were highly resistant to degradation in simulated intestinal fluid for 1 h at 37°C, the longest time tested (Figure 2C, 5C, S6). ssfzd84, a group 2 design with a C18-57 disulfide, showed no discernable degradation (except for the loss of the C-terminal SNAC and his tags) and had an IC50 of 24 (95% CI 14 - 40) pM in the TcdB neutralization assay (Figure 4B).

### Inhibition of the CSPG4 binding site

We applied the same approach to the CSPG4 binding site which was more challenging due to the complex topology of the binding site (Figure 1B). Using the same Rifgen/Rifdock approach followed by ProteinMPNN and sequence optimization (Figure S6), we achieved sub-nanomolar binding against the CSPG4 binding site from two different families of designs, designated group 1 and 2 (Figure 3A,B). Using TcdB2, which preferentially uses CSPG4 for cell entry, we found that these molecules can protect against cytotoxicity with IC50s of 238 (95% CI 43 - 495), 520 (95% CI 85 - 1320) and 332 (95% CI 240 - 444) pM for designs cspg18, cspg27 and cspg35 respectively, after 48 h in the presence of 0.1 pM TcdB (Figure 4C). We stabilized these designs by introducing two disulfide bonds (Figure 3D). After 1 h in the presence of simulated intestinal fluid, intact protein remained compared to complete digestion by 5 minutes for the single disulfide version of the binder (Figure 3D, S8) with no loss in neutralization potency *in vitro* (Figure S8).

**Figure 3.**
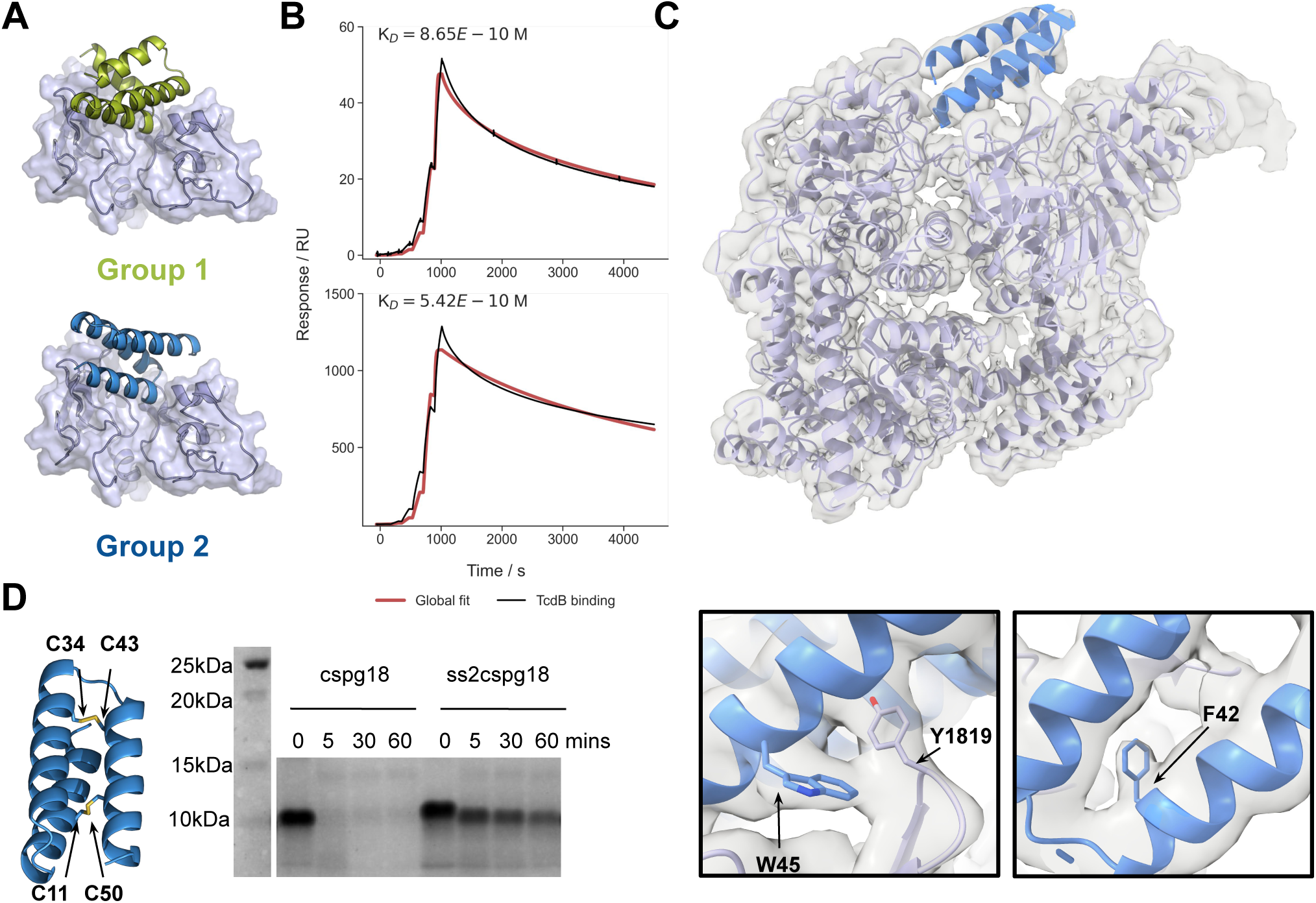
Design of anti-TcdB CSPG4-blocking minibinders. **A.** Design models of the two CSPG4-blocking minibinder families. **B.** SPR traces of a group 1 and group 2 minibinder amine conjugated to the surface with a 4-fold, 6-step dilution series of full length TcdB starting at 100nM. **C.** The design model docked into the CryoEM density map showing agreement between the observed density and the design. Lower inset boxes highlight specific resolved side chains at the target:binder interface. **D.** Disulfide stabilization of CSPG4-blocking minibinders with the location of the two disulfides introduced to make ss2cspg18 (left) and a coomassie stained SDS-PAGE after a time course incubation in simulated intestinal fluid (with 0.1 mg/ml of trypsin and chymotrpysin) at 37°C with cspg18 and the dual disulfide version (ss2cspg18) (right).

We determined the structure of a cspg group 2 binder, cspg67, in complex with TcdB by cryo-EM (Figure 3C). Helices two and three engage with a hydrophobic patch formed from a small two stranded sheet adjacent the native CSPG4 binding cleft (V1816, L1818, Y1819, F1823) and M1831 from a neighboring loop, binding to TcdB in the open conformation. The loop connecting helices 2 and 3 extends into the CSPG4 binding cleft where the C terminus of the second helix forms engages with the polar target surface, primarily through K32 and R36 that flank D1812 on TcdB. Because the binding site is in the central GTD and CPD domains, as opposed to the distal RBD, we were able to resolve bulky side chains (F42, W45 on the binding molecule), which closely align with the design model side chain conformations (Figure 3C).

In order to test the effects of using fzd and cspg minibinders in combination, we tested a titration series of ss2cspg18 (the disulfide stabilized version of cspg18) with 4 concentrations of ssfzd84 (0, 1 pM, 10 pM, 100 pM) in both a CSPG4-dependent set-up using TcdB2 at its EC99 (0.1 pM), and a dual CSPG4-Frizzled-dependent set-up usingTcdB1, also at its EC99 (0.1 pM), on Vero cells expressing both receptors (Figure 4A). The combination of the Frizzled- and CSPG4-blocking minibinders together resulted in an enhanced dose-dependent inhibition of cell toxicity with increasing concentration of ssfzd84, as the curves shift to the left from an IC50 of 520 (95% CI 416 - 671) pM to 35 pM (95% CI 3.7 - 86) pM at 100 pM of ssfzd84 (Figure 4F). There is also an increase of maximum protection to 100%, which is not achieved by ss2cspg18 alone (Figure 4F). These results indicate that combinations of both ss2cspg18 and ssfzd84 provide cooperative protection against TcdB cytotoxicity when both receptors can be used. In the CSPG4-dependent paradigm, there was no additional benefit of increasing ssfzd84 concentrations (Figure 4E).

**Figure 4.**
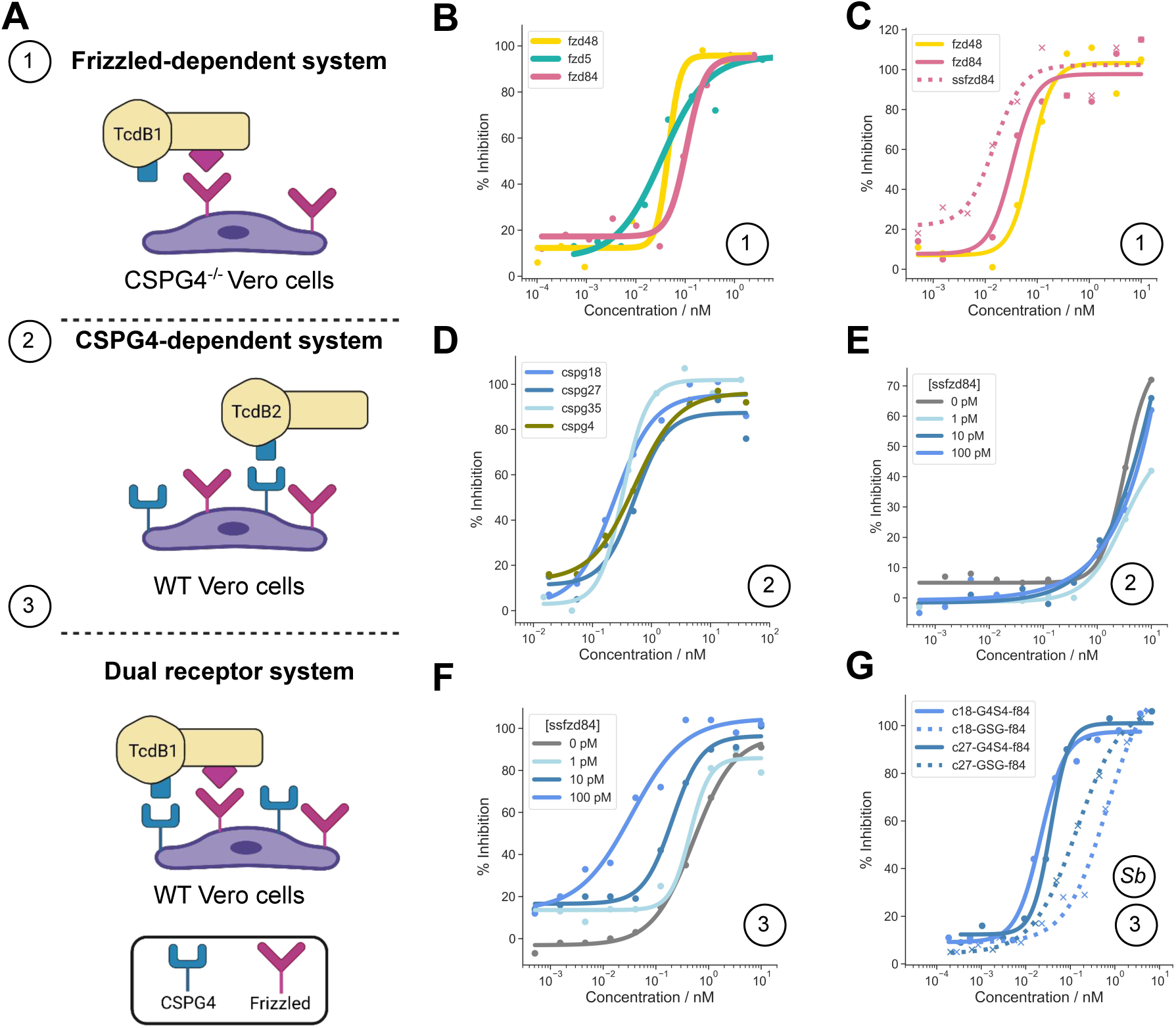
Minibinder neutralization of TcdB in Vero cells. **A.** Overview of the three experimental set-ups used to generate the neutralization data for the different classes of binders alone and in combination. **B.** Sequence optimized fzd binders from group 1 (fzd5) and group 2 (fzd48, fzd84). **C.** Protease resistant, sequence optimized fzd binder ssfzd84 compared with the protease susceptible designs. **D.** Sequence optimized cspg designs from group 1 (cspg4) and group 2 (cspg18, cspg27, cspg35). **E.** Titration of disulfide stabilized ss2cspg18 (x-axis) in the presence of 4 concentrations of ssfzd84 in the CSPG4-dependent system **F.** Titration of ss2cspg18 (x-axis) with four concentrations of ssfzd84 in the dual receptor system. **G.** Neutralization activity of fzd-cspg binder fusions secreted by *S. boulardii*. Fusion constructs between ss2cspg18 and ss2cspg27 fused to ssfzd84 with either a short GSG linker that should not allow simultaneous receptor engagement (dashed) or a longer (G4S)_4_ linker that should enable simultaneous binding at the two sites (solid). In all cases, a single example from independent replicates is shown. Reported IC50 values (in text) are the average across independent replicates. All response curves plotted on the same axes were run in the same experiment.

### Secretion of fzd/cspg fusion constructs in the probiotic yeast *S. boulardii*

Producing biologics *in situ* at the desired site of action is appealing as it would enable continuous production of the biotherapeutic, rather than necessitating repeated oral dosing. *Saccharomyces boulardii* is a probiotic yeast strain that can transiently colonize the lower GI tract(16) that is not susceptible to antibiotic treatment, unlike other experimental bacterial delivery platforms, and has previously been used successfully to deliver antibody-like constructs to the GI tract(17). We therefore tested the production and secretion of minibinders in *S. boulardii*. *In vitro*, the disulfide bond stabilized monomeric minibinders against TcdB were secreted at concentrations up to 300nM (Figure S9). A fusion of the lead CSPG4- and Frizzled-blocking minibinders was also efficiently secreted from *S. boulardii* (Figure S9). Supernatants from *S. boulardii* culture expressing this construct (ss2cspg18 and ssfzd84 fused with a (G4S)x4 linker) neutralized the toxin with an IC50 of 21.8 pM against TcdB1, which can use either CSPG4 or Frizzled to enter cells (Figure 4G). The same fusion linked with only GSG, which is too short to allow simultaneous engagement with both receptor interfaces on TcdB, had a substantially increased IC50 of 672 pM. Thus, avidity achieved by linking binding domains against different regions of the monomeric toxin can enhance activity. These results demonstrate that multiple minibinder formats can be efficiently produced and secreted by *S. boulardii,* setting the stage for miniprotein-based synthetic biotics as therapeutics in a variety of GI disease contexts.

## Discussion

Our designed binders provide a promising new route for combatting the major health challenge presented by the TcdB. Therapeutics that neutralize toxins rather than directly killing pathogens should elicit diminished evolutionary pressure towards antimicrobial resistance, as the toxin is typically not necessary for survival(18). Targeting *C. difficile*-specific virulence factors is particularly appealing because antibiotic-induced disturbances to the microbiota contribute to disease progression and relapse(19). Effective CDI treatment through TcdB neutralization has the potential to improve both clinical outcomes for patients as well as antibiotic stewardship. The possibility of a virulence-factor centered approach has been highlighted by the success of bezlotoxumab, a TcdB targeting monoclonal antibody that reduces the rate of recurrent CDI(2). Despite the clinical benefit of bezlotoxumab treatment, uptake has been limited, largely due to cost(20). Against TcdB, our designed miniproteins target two distinct sites and have very high affinity, protease resistance, and can be secreted from an *S. boulardii* probiotic drug delivery platform to neutralize TcdB. When fused together, they show remarkable potency, in the low picomolar range, in a dual-receptor dependent disease paradigm. The ease of production, either as a recombinant protein or secreted from *S. boulardii*, potential for oral delivery and more targeted approach to CDI control (as opposed to conventional antibiotics), could enable these molecules to be administered prophylactically during *C. difficile* outbreaks to prevent symptom onset or to treat acute CDI at low cost.

The viability of this approach will depend on the ability of these molecules to retain function within the gastrointestinal tract where these proteins will be subject to protease exposure. The stability of these molecules in simulated intestinal fluid containing high concentrations of trypsin and chymotrypsin is promising regarding their ability to neutralize TcdB *in vivo*. Protection studies in a mouse model of CDI using the *S. boulardii* platform to secrete the most promising fusion constructs are currently being pursued. Neutralizing minibinders that target other infectious disease-associated toxins, particularly those that drive gastrointestinal disease, may be readily designed, and the concept extended to other features of infection biology, such as pathogen biofilms and inflammatory cytokines.

## Materials and Methods

### Design and optimization of Frizzled- and CSPG4-blocking minibinders

Docks of both Frizzled- and CSPG4-blocking minibinders were generated using a Rifgen-Patchdock-Rifdock procedure as described in depth previously(7, 21, 22). The input structure for Frizzled-blockers was 6C0B(11) with the Frizzled chain removed and the model cropped to remove the distal arms that were far from the Frizzled binding site (for computational efficiency). The docking procedure used a library of 30,000 helical bundles that were docked against hotspot residues M1437, L1483, L1488, V1491, Y1509 and F1597. For CSPG4-blocking binders, the input PDB used was 7ML7(13), again with the receptor chain removed. The hotspot residues selected for the docking procedure were V1816, L1818, Y1819, F1823 and M1831. The Rifdock outputs then underwent sequence design using ProteinMPNN as described above with one sequence generated prior to FastRelax and one post FastRelax at temperature 0.001. For Frizzled blocking binders, a pLDDT cutoff of 90 was used alongside a PAE interaction of < 8 to reach 15,000 designs. For CSPG4, only 2811 designs were ordered, all of which had a PAE interaction < 10.

Initial hits were validated for binding via BLI and TcdB neutralization assays (data not shown) and then underwent a site saturation mutagenesis screen via yeast surface display (described below). The most promising mutations for each design on the basis of affinity increase without making the binding surface substantially more hydrophobic were selected to be included in a combination library which was ordered as an IDT ultramer with approximately 1E6 diversity. The combo libraries were assembled from two ultramers via PCR extension where necessary due to length constrictions and then cloned into yeast as described below. The sequences for the combo libraries are included in Table S2.

### Disulfide stabilization of fzd and cspg designs

Disulfides were introduced using a previously published disulfide stapler(23) that can identify sites which accommodate disulfide bonds through searching a database of 30,000 examples of disulfide bonds in native proteins. A single site was identified in each of the backbones for group 1 and group 2 fzd binders between residues 28 and 44 in group 1 and 18 and 57 in group 2. Within the group 2 cspg designs, possible disulfide sites were between residues 11 and 50, 23 and 53, and 34 and 43. Two pairs of dilsulfide bonds were tested in each of the neutralizing designs. In each case, the disulfide stabilized designs were expressed at the 4 mL scale, as described below in the *E. coli* expression section. Fzd designs expressed well in BL21 (NEB) whereas cspg designs were expressed in T7 shuffle express (NEB).

### Electron microscopy of TcdB complexes

#### CryoEM of TcdB:fzd48

TcdB was thawed and complexed with 3 fold molar excess minibinder fzd48 at room temperature for 10 min before dilution to 0.81 mg/mL TcdB concentration in buffer (Tris/HCl pH 7.5 (25 mM) NaCl (150 mM)). Samples applied onto freshly glow-discharged grids (-15 mA, 25 s, 0.39 mBar) 2/2 C-flat holey carbon grids in the chamber of a Mark IV Vitrobot, with humidity held at at 100% and temperature at 22°C. The wait time before blotting was 7.5s, with a blot time of 7.0 s and a blot force of 0, after which the grid was plunge frozen in liquid ethane.

Cryo-EM grids were screened and data was collected on a ThermoFisher Titan Krios transmission electron microscope (FEI Thermo Scientific, Hillsboro, OR) operated at 300 kV and equipped with a Selectis energy filter and Gatan K3 direct detector. Data collection was automated with SerialEM(24). A total of 6713 movies were collected: 2,186 with no stage tilt, 3,873 at a 15° stage tilt, and 654 at a 30° stage tilt. Movies were acquired in super resolution mode at a nominal magnification of 105,000x (0.4215 Å/pixel super-resolution pixel), fractionated in 100 frames at 9.4 e^-^/Å^2^ /sec for a total exposure of ∼47 e^-^/Å^2^ over 5.0 seconds. All data processing was carried out in cryoSPARC v4.4. Alignment of movie frames was performed using Patch Motion Correction(25). Defocus and astigmatism values were estimated using Patch CTF with default parameters.

Curate Exposures was used to remove 750 micrographs, leaving 5963 good micrographs. 1,068,213 particle images were initially selected using Blob Picker with a minimum and maximum particle diameter of 200 and 400 Å extracted in 920 x 920 pixel boxes and Fourier cropped to 460 x 460 pixels. Following 2D classification the five best classes, containing 39,344 particle images, were used to train Topaz on all 5963 good micrographs. Topaz extract was used to select 2,755,304 particle images, extracted in 920 x 920 pixel boxes and Fourier cropped to 460 x 460 pixels. This procedure was followed by 2D classification with 300 classes, 86 iterations 11 of which were final iterations using all particles. The 15 best classes, containing a total of 148,808 particle images, were subjected to 2D classification again using 50 classes. The best 23 classes, containing a total of 108,076 particle images, were used for 3D *ab initio* structure determination with C1 symmetry. The initial structure was refined with Non Uniform Refinement(26) with C1 symmetry for a final global resolution estimate of 4.6 Å, including correction for the effects of masking. The final map was sharpened using DeepEMhancer(27). 3D maps for the half maps, final unsharpened maps, and the final sharpened maps were deposited in the EMDB under accession number ####. Local resolution estimation was run on the final Non Uniform Refinement map.

The published crystal structure of TcdB (pdb ID: 6OQ5)(28) was used as an initial reference for building the final cryoEM structure. The model was fit into density using UCSF ChimeraX(29). Isolde(30) in UCSF ChimeraX and Coot(31) was used to better fit portions of the model to the map. Phenix(32) was used to trim the model to polyA, before further refinement in Coot and Isolde. Phenix and MolProbity(33) were used assess model quality throughout. The de novo minibinder fzd48 was added last to remove bias, with further refinement in Coot and Isolde. Figures were generated using UCSF ChimeraX. The final structure was deposited in the PDB under number ####.

#### CryoEM of TcdB:cspg67

For the CryoEM of TcdB and CSPG4-blockers, cspg67 belonging to the group 2 family of cspg designs was used, which is the same dock (ie both group 2 designs) as the lead candidates ss2cspg18. Purified TcdB in a buffer containing 50 mM NaCl and 20 mM tris at pH 8 was mixed with CSPG4 minibinder #67 at a 1:1 molar ratio and incubated on ice until use. Three microlitres of the mixture was applied to nanofabricated holey gold grids(34) which were previously glow discharged in 40 mbar air for 40 s with 25 mA current. The sample was applied in the environmental chamber of a Leica EM GP2 plunge freezing device (Leica Microsystems) at 4°C and ∼80% humidity, with a 30 s preincubation time and a 2-3 s blot time before plunge freezing in liquid ethane.

The specimen was imaged with a Titan Krios G3 300 kV cryo-electron microscope equipped with a Falcon 4i direct detector (ThermoFisher Scientific). The specimen was imaged at a nominal magnification of 75,000x, corresponding to a calibrated pixel size of 1.03 Å. Data collection was automated with the EPU software (ThermoFisher Scientific) and was monitored with cryoSPARC Live. A dataset consisting of 8402 movies was acquired with 0° stage tilt. A second dataset consisting of 4288 movies was acquired with 30° stage tilt. Movies were acquired at ∼6.8 e/Å^2^/s for a total exposure of ∼42 e/Å^2^. The resulting movies were stored in the EER file format(35).

All cryoEM data were processed with cryoSPARC v4.4.0(36). Initial patch-motion correction and patch-CTF estimation were carried out during data collection using cryoSPARC Live. Raw movie frames were grouped into 40 fractions with the upsampling factor set to 1 (i.e. no superresolution information used). Particle images were initially selected from a subset of the dataset using blob picker in cryoSPARC. After several rounds of 2D classification, a curated particle dataset was used for training a neural network model using the Topaz package(37). The Topaz model was then used for selecting particle images from all micrographs. Particle images were classified with iterative rounds of 2D classification and heterogenous refinement. Particle images of TcdB:cspg67 complexes were separated from the unbound TcdB particle images by 3D classification using a mask encompassing the cspg67 minibinder volume. The downstream cryo-EM refinements were carried out with the TcdB:cspg67 particle images only.

Reference-based motion correction and global CTF refinement were carried out to improve the resolution of the final map. The final non-uniform refinement map reached a resolution of 3.0 Å. The cryo-EM maps were deposited in the EMDB under accession number ####.

Initial atomic model building was carried out by docking an *apo* TcdB atomic model built in a previous cryo-EM study and originally derived from a predicted alphaFold model (Uniprot entry: P18177; Miletic et al., in preparation). A predicted model for cspg67 was docked into the TcdB:cspg67 complex cryo-EM map and combined with the TcdB model using ChimeraX(29). The model was then manually adjusted with ISOLDE(30) and refined with phenix.real_space_refine(38). The atomic model of the TcdB:cspg67 complex was deposited in the PDB with accession code #####.

### Yeast surface display

Both the initial design library and subsequent SSM libraries for the Frizzled-blocking binders (fzd library) and CSPG4-blocking binders (cspg library) were ordered as chip DNA oligos (Agilent). The initial library size was 15,000 and 2811 for the fzd and cspg libraries, respectively. The fzd designs were flanked with 5’ GGTGGATCAGGAGGTTCG and 3’ GGAAGCGGTGGAAGTGGG adapters and cspg had 5’ TCGTCTGGTAGTTCAGGC and 3’ GGTTCTAGTGGCTCATCG adapters to enable subpool amplification. Libraries were qPCR amplified using Kapa hifi polymerase (Kapa Biosystems) in duplicate 25 µL reactions with the first reaction used to determine the cycle number to achieve half maximal amplification. Then a second production run was performed with these reactions purified from an agarose gel. Lastly, the gel purified DNA fragments were amplified again to obtain sufficient material for electroporation in EBY100 with 2 µg of linearized petcon3 and 6ug of insert(39).

Yeast cultures were maintained in C-Trp-Ura media with 2% glucose (CTUG) at 30°C. 18 h prior to sorting, 10 mL of SGCAA with 0.2% glucose was inoculated with 250 µL of C-Trp-Ura culture. In preparation for sorting, the cells were harvested at 4000 x g for 3 min followed by a wash step in PBS with 1% BSA (PBSF). Both yeast libraries and SSMs underwent four sorts. We began with an expression sort staining with only anti-myc FITC (Immunology Consultants Laboratory) followed by two rounds of ‘avidity sorts’ where the cells were stained with both anti-myc FITC and streptavidin conjugated to phycoerythrin (SAPE, Thermo Fisher) mixed with target protein in a 1:4 ratio. For the fzd library, the biotinylated RBD toxin fragment was used while for the cspg library the biotinylated TcdB 1-2100 toxin was used. Finally we finished with a titration sort with streptavidin-PE conjugated to target protein in a 1:1 ratio to rank the binders by SC50, as described previously(7). For each sort, cells were collected then grown overnight in CTUG. 1 mL of culture was harvested and plasmid DNA extracted using a yeast plasmid miniprep kit (Zymo) and prepared for illumina sequencing through two rounds of PCR to attach pool specific barcodes. Reads were assembled using PEAR(40) and SC50s determined using a set of previously published computational tools from Cao *et al*(*7*). For combo libraries, no SC50s were calculated and rather the most represented designs at the lowest sort concentration were taken forward. For fzd designs, the top 94 based on abundance at 37 pM sorting concentration were taken forward for affinity screening. For cspg binders, due to the difficulty of working with full length TcdB on surface based kinetic assays, the top 24 designs ranked by abundance at 20 pM were tested in neutralization assays. The titration series for the libraries are included in Table S1.

### TcdB expression

Plasmid pHis1522 encoding his-tagged TcdB1 (VP10463) was a kind gift from Hanping Feng (University of Maryland Dental School, Baltimore, MD, 21201, USA.), and the plasmid for TcdB 027 in pHis1522 was synthesized by Genscript USA. TcdB was purified from *Bacillus megaterium* (MoBiTec, Germany) carrying the vector pHIS1522 encoding C*. difficile* TcdB VP10463 or 027 fused to a C-terminal His tag. From glycerol stocks, overnight starter cultures of *B. megaterium* were grown in lysogeny broth (LB) with tetracycline selection. The next morning, starter cultures were used to inoculate 1-2 L of terrific broth (TB) with tetracycline selection, and were grown at 37°C, 180 RPM until the OD600 reached 0.8 or higher, upon which xylose (0.5% w/v final concentration) was added to induce expression of TcdB overnight at 30°C, 180 RPM. The following morning, cells were pelleted at 4000 x g for 12 min and pellets were resuspended in lysis buffer containing 20 mM Tris pH8, 0.1 M NaCl, 1 mg/ml lysozyme, 1% v/v protease inhibitor cocktail P8849 (MilliporeSigma), and 100 U/ml Pierce universal nuclease (88701, ThermoFisher). The suspension was then passed twice through an EmulsiFlex C3 microfluidizer (Avestin) at 15,000 psi to lyse the cells. Cell lysate was clarified by centrifugation at 14,000 × g, 4°C for 20 min and supernatants were filtered with a 0.2 µM filter prior to running on a 5 ml HisTrap FF Ni-NTA column using an Äkta fast protein liquid chromatography (FPLC) systems (Cytiva). Proteins were eluted using a gradient of buffer containing 500 mM imidazole and a single peak was collected, and then further purified on a 1 ml HiTrap Q column (Cytiva). Proteins were eluted using a gradient of buffer with 100 mM to 1 M NaCl. When required for cryoEM purposes, collected fractions containing TcdB were then run by size exclusion on a Superose 6 increase 10/300 GL (Cytiva) to further polish the protein preparation and to buffer exchange TcdB into 20 mM Tris, 50mM NaCl pH 8.0. Purified protein was either directly used for cryoEM analysis, or flash frozen in liquid nitrogen with 10% glycerol and stored at -80°C.

For yeast screening purposes, C-terminal truncated TcdB1 (amino acids 1-2100) with inactivating W102A/D286N/D288N mutations in the glucose transferase domain was generated in the pHis1522 vector, and purified protein was isolated as described above. Biotinylation was performed by incubating purified TcdB with a 15-fold molar excess of maleimide biotin (ThermoFisher) overnight at 4°C in 20mM Tris pH7.5, 150mM NaCl, 5% glycerol. The biotinylated TcdB was then purified on a 1 ml HiTrap Q column (Cytiva).

Alternatively, TcdB 1285-1804 with C-terminal Avitag (RBD toxin fragment) was generated and purified as above, followed by overnight biotinylation reaction at 4°C using a BirA biotin labeling kit (Avidity), then desalting with a Zeba 40k MW biotin-removal spin column (ThermoFisher). Successful biotinylation was verified by Western blot, using streptavidin-horse radish peroxidase as the detection agent.

### *E. coli* expression of designed proteins

Designed proteins were ordered as eblock fragments (Integrated DNA Technologies) with flanking BsaI cut sites and then cloned into LM0627 (addgene 191551) which encodes an N-terminal MSG and C-terminal SNAC(41) and 6xhis tag. An Echo acoustic liquid handler (Beckman Coulter) was used to dispense 1 µL reaction volumes (0.1 µL water, 0.1 µL T4 ligase buffer, 0.375 µL eblock fragment at 4 ng/µl, 0.06 µL of BsaI-HFv2, 0.1 µL T4 ligase, 0.275 µL destination vector at 50 ng/µl). These reactions were incubated at 37°C for 20 min before inactivation at 60°C for 5 min. 5 µL of BL21 competent cells (or, in the case of the disulfide stabilized cspg binders, T7 shuffle express) (NEB) were dispensed onto the 1 µL reactions, and then incubated on ice for 30 min before 10 s heat shock at 42°C. After 1 h recovery at 37°C in 100 µL SOC media, 4×1 mL cultures of autoinduction media were inoculated with 25 µL of transformed cells. These were grown for 20 h at 37°C.

Cells were harvested through centrifugation at 4000 x g for 5 min and then lysed with 100 µL of BPER supplemented with 0.1 mg/ml lysozyme, 10 µg/ml DNAse I and 1 mM PMSF. The pellets were incubated in lysis buffer for 15 min on a shaker at 1000 RPM before pooling and clarifying the lysates via centrifugation at 4000 x g for 10 min. The supernatant was then applied to 50 µL of Ni-NTA resin (Thermo Fisher), washed 3x with 300 µL of wash buffer (20mM Tris, 300mM NaCl, 25mM imidazole) and then eluted in 200 µL of elution buffer (20mM Tris, 300mM NaCl, 500mM imidazole pH 8). Elutes were injected onto a S75 5-150 GL Increase column via autosampler and underwent size exclusion chromatography into PBS (or HBS-EP for SPR experiments as described below). Fractions were normalized to 10 µM using an OT2 (Opentrons). Aggregation states were determined by plotting the elution volume of each peak relative to a standard curve for that column and yields were determined via integration of the SEC elution curve adjusted for 4 mL culture volume.

### Surface plasmon resonance

For all SPR experiments, the analyte proteins (that which is flowed over the chip) were purified in HBS-EP (0.01 M HEPES pH 7.4, 0.15 M NaCl, 3 mM EDTA, 0.005% v/v Surfactant P20) (Cytiva). All experiments were run at 25°C. For both the fzd binders, approximately 500 units of biotinylated RBD of the respective toxin was captured onto a biotin capture chip (Cytiva). For the single cycle kinetic runs, the binders were injected at 30 µL/min for 120 s, followed by 60 s dissociation and injection of the next concentration for six steps. The concentration series used in each experiment is described in the associated figure legend. After the sixth step, the dissociation was measured for 10 min to > 1 h (as indicated on the figure axis). For cspg binders where the full length toxin had to be used, the miniprotein was captured on a CM5 chip through amine conjugation (Cytiva). Then, the full length toxin in HBS-EP was flowed over in a six step single cycle kinetics run, in the same fashion as for the RBD binders. All single cycle kinetic data is shown with the measured data in black and the global Langmuir 1:1 fitting in red. The fitting was done using the Biacore 8K Evaluation software (Cytiva) and then replotted in seaborn.

### Simulated intestinal fluid assay

For both simulated intestinal fluid (0.1 mg/ml trypsin, 0.1 mg/ml chymotrypsin, 3 mM sodium taurocholate, 19 mM maleic acid, 34.8 mM NaOH, 68.6 mM NaCl and 0.2 mM lecithin pH 6.5) and simulated gastric fluid (45 mM NaCl, 100 mM HCl, 2 mg/ml pepsin), 25 µM of binder was incubated at 37°C for 5, 10 or 60 min (0 minute condition was without protease). The reaction was quenched through the addition of 1 mM PMSF followed by incubation at 95°C for five min. These samples underwent SDS-PAGE using a Criterion TGX stain free gel (BioRad) at 200V for 30 min before being stained with coomassie blue.

### TcdB neutralization assays

Vero (ATCC) and Vero CSPG4-knockout cells(42) were grown in DMEM with 10% fetal bovine serum in the presence of penicillin and streptomycin, at 37°C, 5% CO2.

For cell assays that test TcdB toxicity through both the FZD and CSPG4 receptors, Vero cells were seeded at 5000/well in 96 well clear CellBind plates (Corning) and used 24 hours post plating. Test miniproteins were serial diluted in PBS, then added using a Bravo liquid handler (Agilent), followed immediately by recombinant TcdB1 to a final concentration of 0.1pM. Cell viability was assessed 48 hours later, when >99% of the vehicle control cells appeared to be rounded, by adding alamarBlue (ThermoFisher) and reading fluorescence signal (ex:555; em:585) 3 hours later in a Spectramax m5 plate reader (Molecular Devices).

Alternatively, for cell assays favoring TcdB entry through FZD receptors, Vero CSPG4-knockout cells were seeded in 96 well plates at 5500 cells/well. Using a Bravo liquid handler (Agilent), serial diluted test samples were added to cells, immediately followed by TcdB1 to a final concentration of 1.5 - 3 pM. Viability was assayed by adding alamarBlue Cell Viability Reagent (aka Resazurin, Thermofisher) after microscopic examination indicated cell rounding in the vehicle control wells (up to 48 hours post TcdB addition), and reading fluorescence signal (ex:555; em:585) 3 hours later in a Spectramax m5 plate reader (Molecular Devices).

For testing cspg minibinders, recombinant TcdB2 was used because it binds preferentially to the CSPG4 receptor. Vero cells were seeded at 5000/well in 96 well plates, and incubated overnight. Serial diluted samples were added to wells using a Bravo liquid handling robot (Agilent), immediately followed by TcdB 027 to a final concentration of 0.1 pM. Cell viability was assessed after 24 hours by adding alamarBlue Cell Viability Reagent (aka Resazurin, ThermoFisher), when microscopic examination indicated cell intoxication (ie., cell rounding, apoptotic bodies) in the vehicle control cells, and reading fluorescence signal (ex:555; em:585) 3 hours later in a Spectramax m5 plate reader (Molecular Devices).

Neutralization curves are a representative curve from independent replicates while the reported IC50s are the mean from across replicates with the 95% confidence interval.

### Secretion of minibinders from *S. boulardii*

Minibinder fusions to the secretion signal of the Mat-alpha mating pheromone from *Saccharomyces cerevisiae* were expressed in a *S. boulardii* MYA-796 wild type strain using the *ADH2* constitutive promoter in a 2 micron plasmid vector. The probiotic genome was not further engineered to increase secretion levels in order to avoid potential impacts on yeast fitness and concomitant effects on yeast survival in the GI tract In particular, while overexpression of protein disulfide isomerase can improve the secretion of proteins containing disulfide bonds, such as single-chain antibody fragments, we found that miniproteins with engineered disulfide bonds expressed well in a wild type context and did not compromise yeast fitness, consistent with their successful expression in BL21/T7 shuffle express.

*S. boulardii* cells were grown in liquid XY media (2% bactopeptone, 1% yeast extract, 0.01% adenine sulfate, 0.02% tryptophan), 200 ug/mL G418, 2% ethanol, 2% glycerol for 3 days at 30°C. Cells were centrifuged and supernatants were were pelleted by centrifugation and concentrated 10-fold with a speedvac (Vacufuge) at 45°C. Proteins were separated by 20% acrylamide/2% bisacrylamide SDS-PAGE. Minibinders in the supernatant were quantified with a direct ELISA using the nanobody Sb68-FLAG as a reference for the standard curve. For ELISA, the primary FLAG antibody used was conjugated with HRP (Sigma, A8592-1MG).

#### Data Visualization

Figures were produced using PyMol, ChimeraX(29), Seaborn(43) and biorender.com.

## Acknowledgments

We thank Stephanie Berger, Ingrid Swanson Pultz, Basile Wicky, Lukas Milles for helpful discussions; Kandise VanWormer, Hernan Nunez-Ortega, Rafael Ticzon, and Andre Dubief for lab support; Luki Goldschmidt and Patrick Vecchiato for technical support; Dionne Vafaedos and Nicole Roullier for yeast transformation and NGS library preparation; Joel Quispe and Sasha Dickinson for management of the Titan Krios at the University of Washington; Alexis Courbet for assistance in Topaz configuration at UW.

R.J.R is a Washington Research Foundation Postdoctoral Fellow. H.L. is a recipient of the Pfizer Graduate Scholarship. This work was supported by the Howard Hughes Medical Institute grant number 0001091096 (D.B.), the Department of the Defense, Defense Threat Reduction Agency grant HDTRA1-21-1-0007 (M.G. and R.J.R.), Howard Hughes Medical Institute C19 HHMI INITIATIVE (M.G., R.J.R.), The Audacious Project at the Institute for Protein Design, the Bill & Melinda Gates Foundation #INV-010680, NSERC Discovery Grant, and Canadian Institutes of Health Research Project Grant (R.A.M). CryoEM data for the TcdB:cspg67 sample was collected at the Toronto High-Resolution High-Throughput cryoEM facility, supported by the Canada Foundation for Innovation and Ontario Research Fund.

## Author Contributions

R.J.R., R.A.M., and D.B. conceived of the study. R.J.R. designed the proteins, did the yeast surface display experiments, *E. coli* expression of designs, binding screens, disulfide stabilization and protease stability assays. J.T. did the TcdB neutralization assays supported by H.L.. R.P. developed the *S. boulardii* platform, produced the *S. boulardii* strains and characterized the secreted protein product. S.M. determined the structure of the CSPG4-blocking minibinder bound to TcdB with support from Z.L and J.L.R. C.W. determined the structure of the Frizzled-blocking minibinder with support from K.D.C and A.J.B. M.G., X.W. and G.L.B. advised on experiments. B.C. advised on design considerations. J.L.R., D.S., M.T., R.A.M., and D.B. were responsible for funding acquisition and supervision of this work. R.J.R. and D.B. wrote the first draft of the manuscript. All authors read and contributed to the manuscript.

---

**Figure S1.**
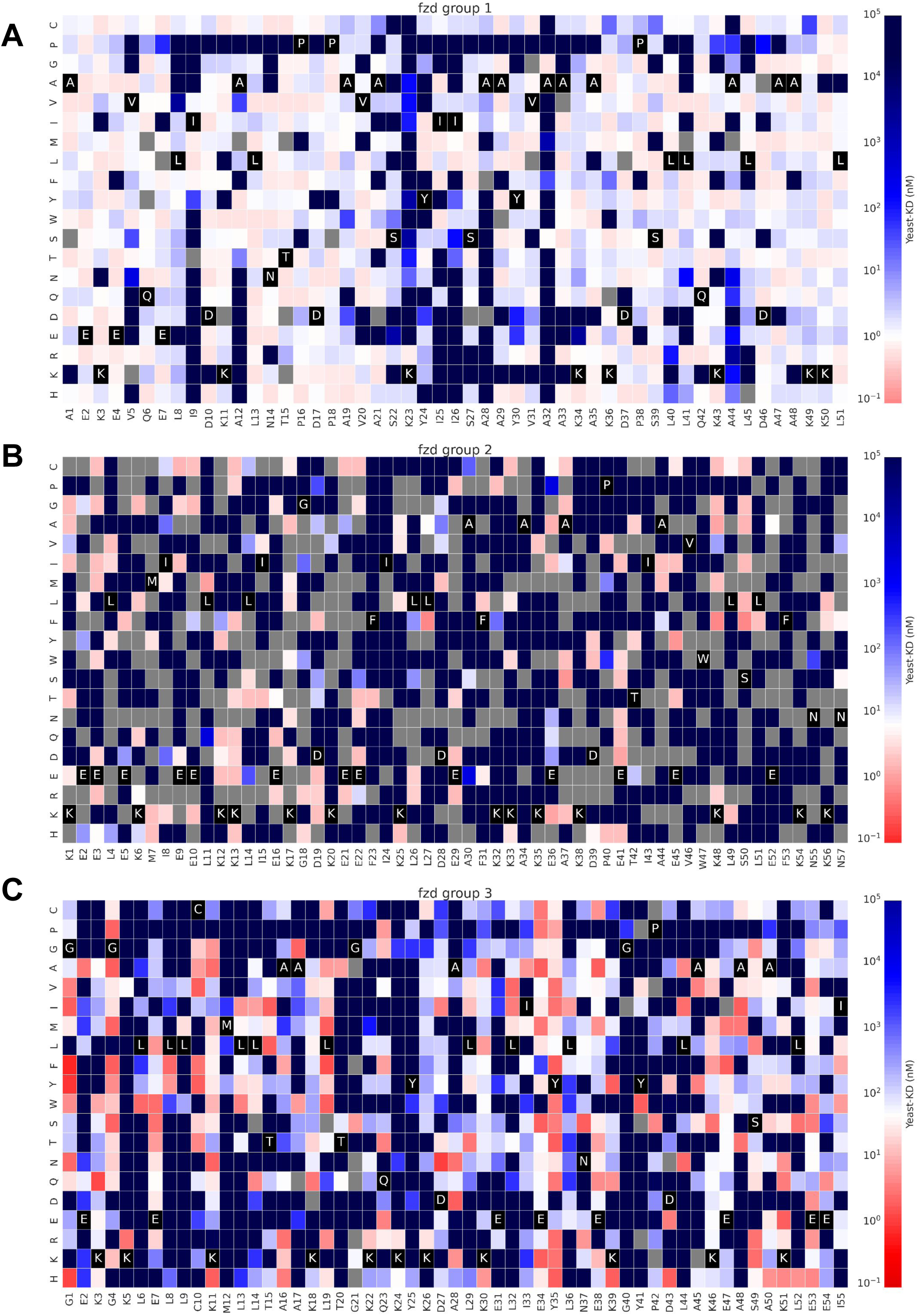
Yeast surface display SSM of group 1 (A), group 2 (B) and group 3 (C) fzd parental designs from which all the sequence optimized variants are derived. Yeast KD is the SC50 as defined by Cao *et al.* (2022). Gray squares indicate that the variant was not identified in the library. Group 2 has poor library coverage, but was still taken forward as it was among the most promising initial hits based on preliminary neutralization data.

**Figure S2.**
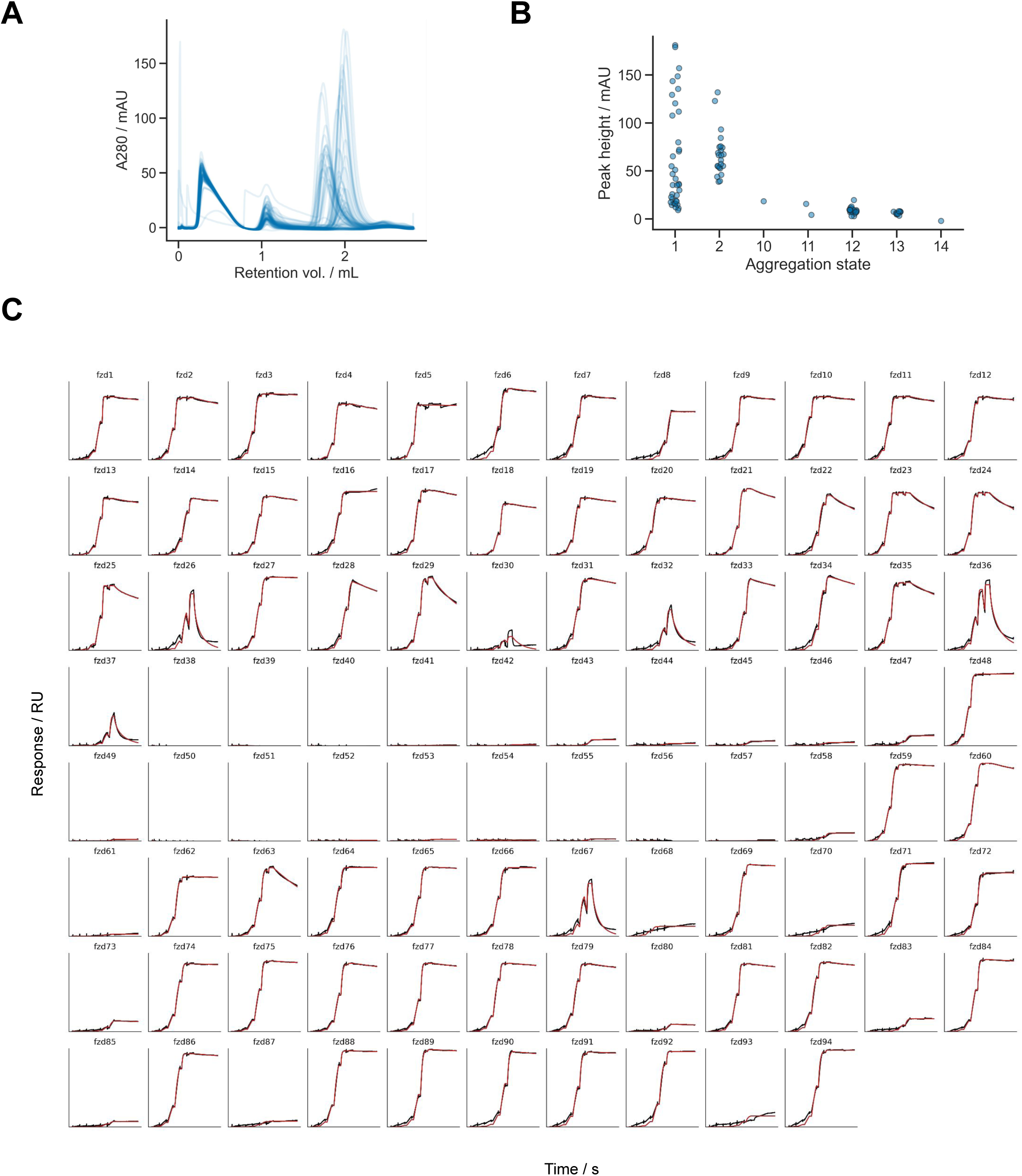
Optimization of Frizzled-blocking miniproteins. **A.** SEC traces of 94 designs sequence optimized designs from 4 mL culture. **B.** Aggregation state of each design based on SEC profile and molecular weight standard curve for the column. **C.** Affinity determination through SPR with the RBD of TcdB captured on the chip and a 6-step 5-fold dilution series of each miniprotein starting at 25 nM. Designs fzd38 to fzd47 and fzd49 - fzd58 failed to express, rather than failed to bind. Global fit is shown in red while the measured data is shown in black.

**Figure S3.**
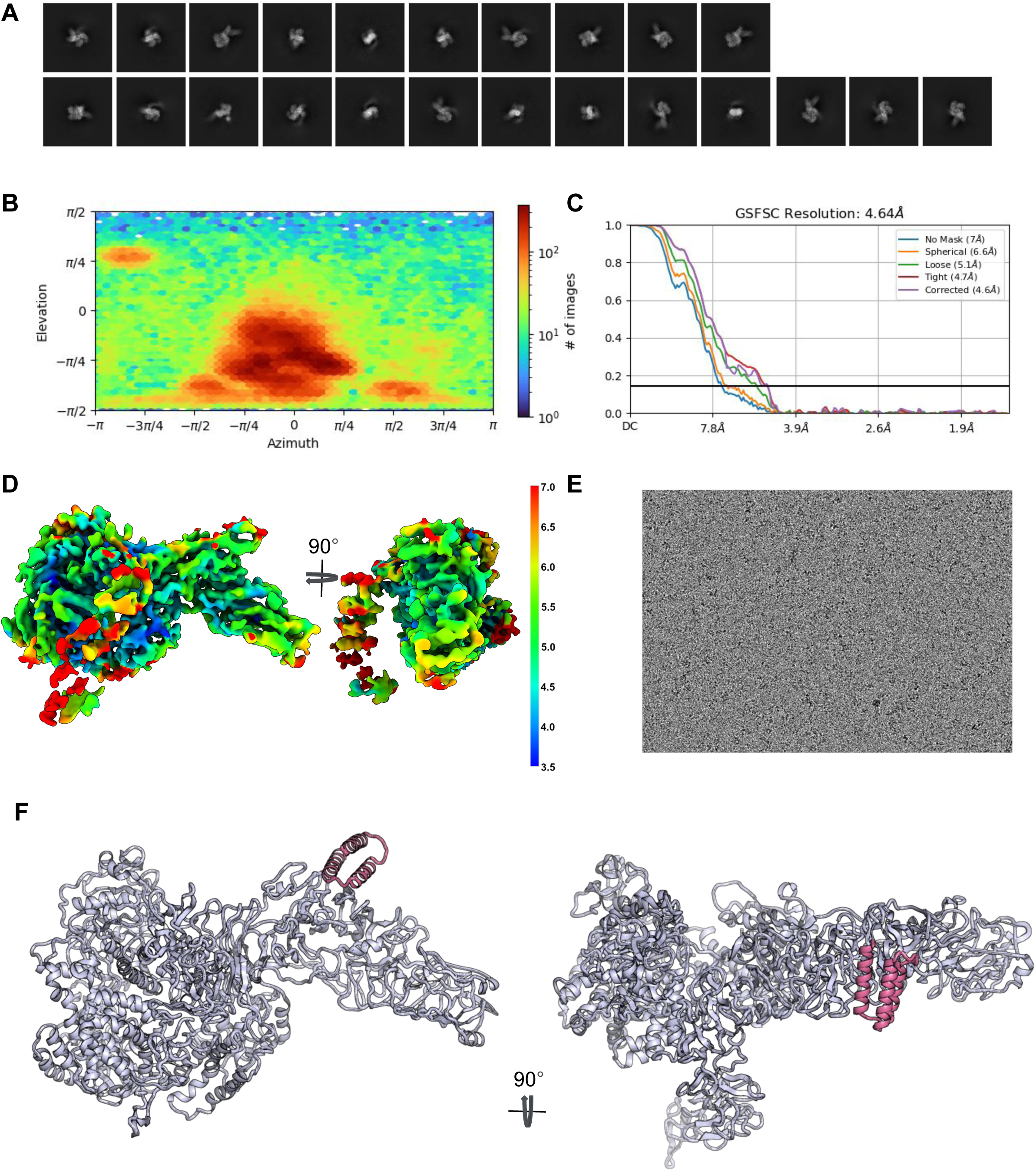
Determination of cryoEM structure of fzd48 bound to TcdB. **A.** Representative 2D class averages. **B.** Orientational distribution plot. **C**. Global Fourier Shell Correlation (FSC) following a gold standard refinement and with correction for the effects of masking. **D**. CryoEM density colored by resolution indicated by the scale bar (unit is Å). E. Representative micrograph. **F.** CryoEM model of fzd48 bound to TcdB.

**Figure S4.**
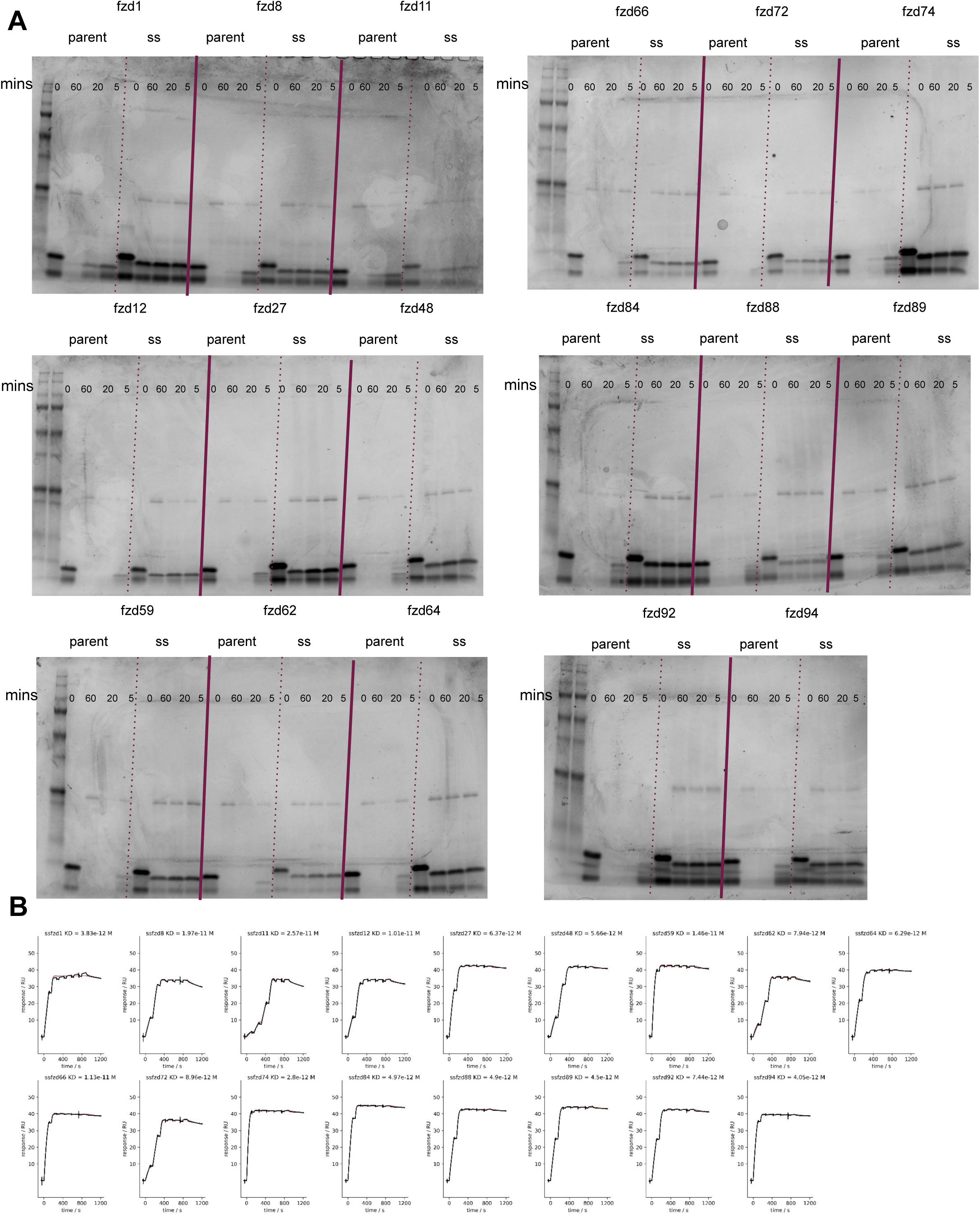
Disulfide stabilization of the Frizzled-blocking minibinders from group 1 and group 2. **A.** SIF assay where designs are incubated in simulated intestinal fluid for 5, 20 or 60 min. The 0 time point contains no protease. **B.** SPR traces of disulfide stabilized designs across a 6-step 2-fold dilution series starting at 50 nM. Global fit is shown in red while the measured data is shown in black.

**Figure S5.**
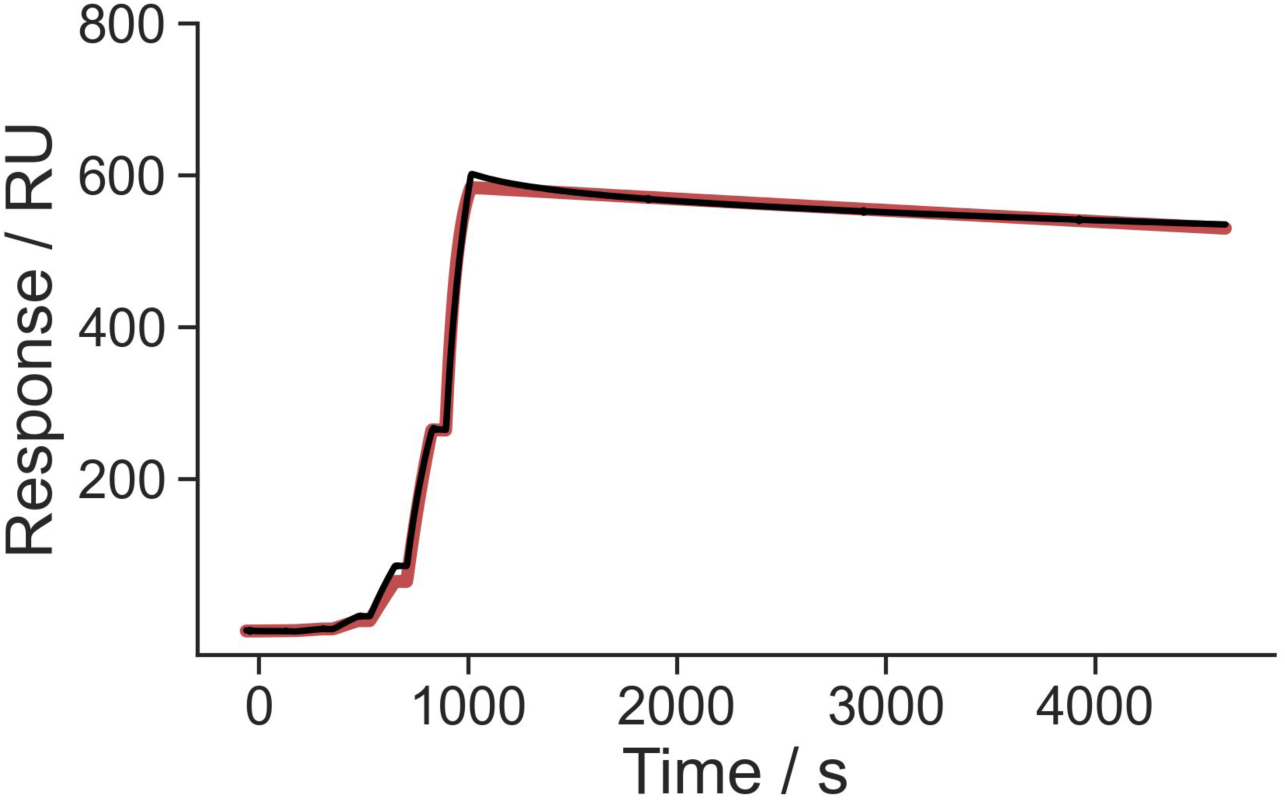
Binding of ssfzd84 to full length TcdB. ssfzd84 was immobilized via amine conjugation. Full length TcdB was used as an analyte over a 6-step 4-fold dilution starting at 100 nM.

**Figure S6.**
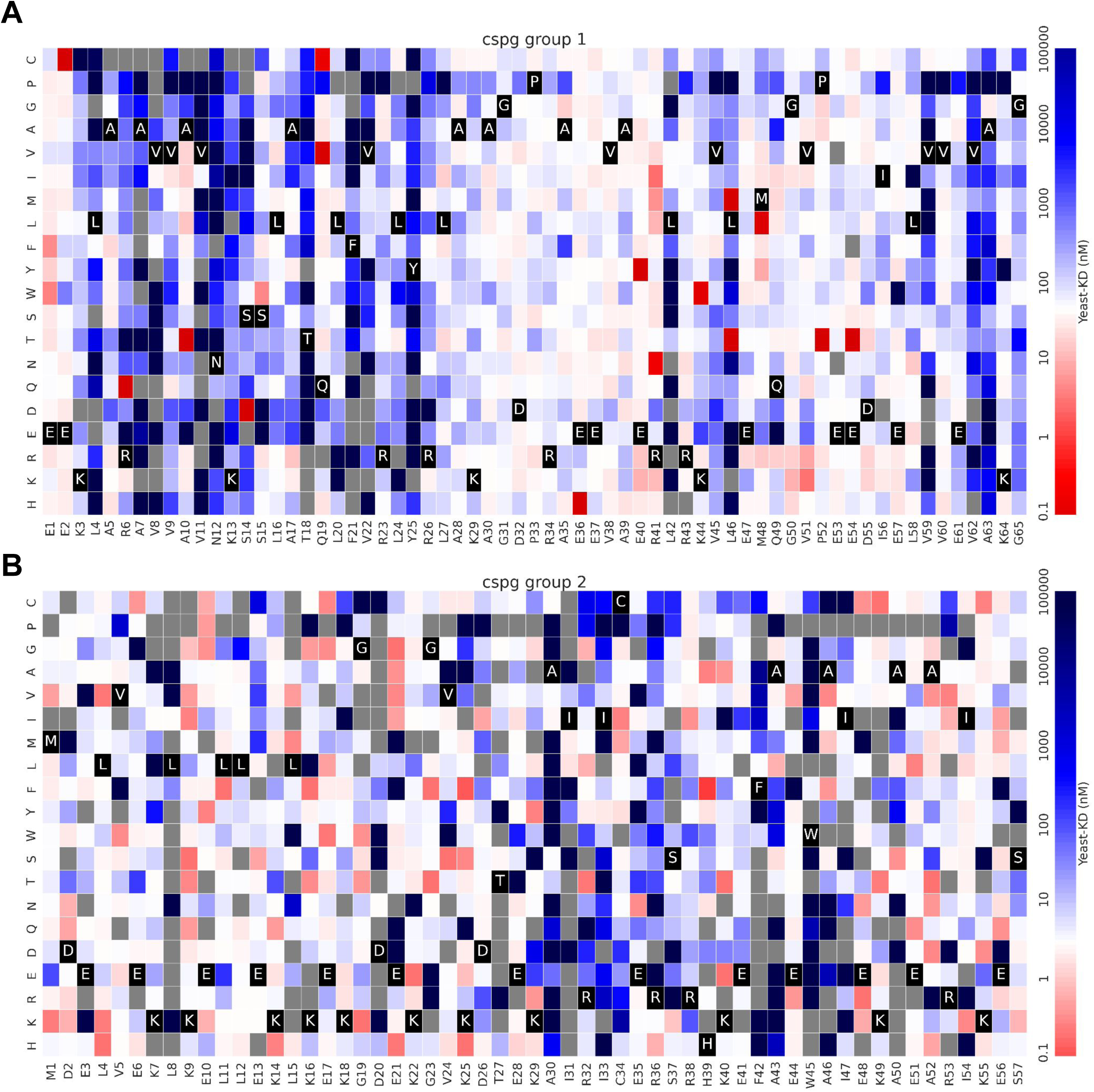
Yeast surface display SSM of group 1 (A) and group 2 (B) cspg parental designs from which all the sequence optimized variants are derived. Yeast K_D_ is the SC50 as defined by Cao *et al*. Gray squares indicate that the variant was not identified in the library.

**Figure S7.**
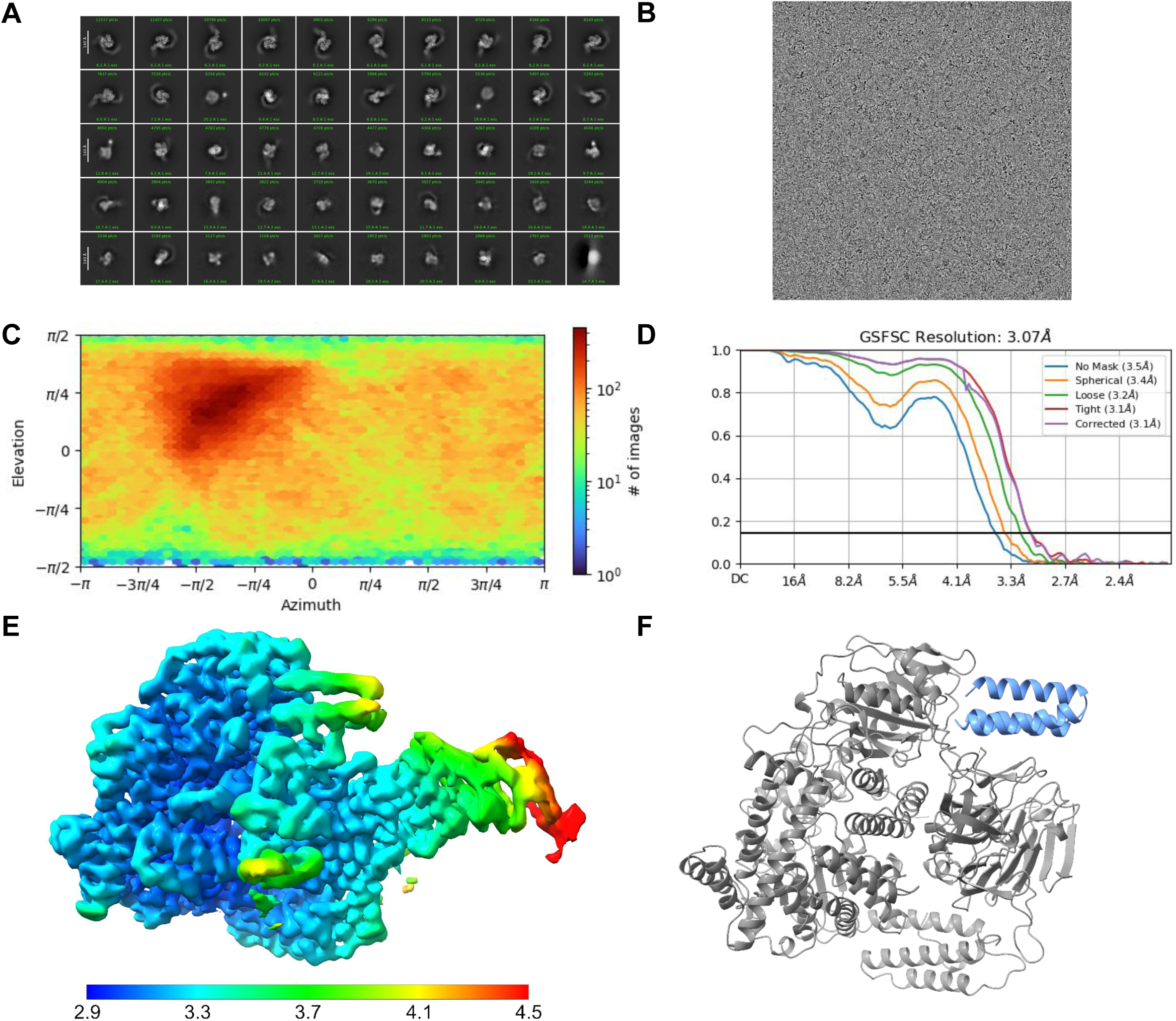
CryoEM structure of TcdB:cspg67. **A**. 2-D class averages **B.** Representative micrograph **C.** Orientational distribution plot. **D.** Global Fourier Shell Correlation (FSC) following a gold standard refinement and with correction for the effects of masking D. CryoEM density colored by resolution indicated by the scale bar (unit is Å) **E.** CryoEM model of cspg67 bound to TcdB.

**Figure S8.**
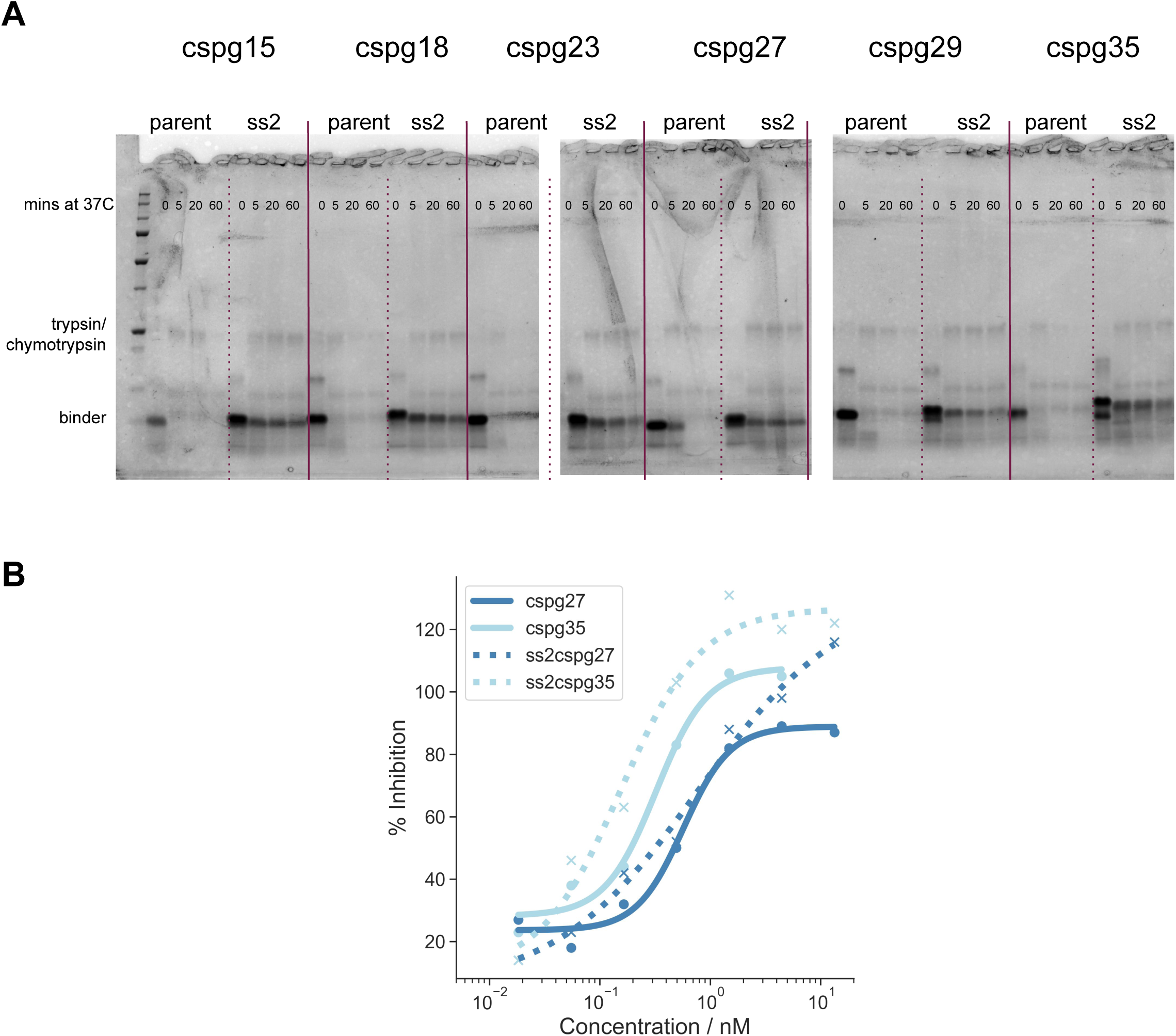
Introduction of two disulfide bonds to enhance protease stability. **A.** Time course incubation in simulated intestinal fluid at 37°C for the parental designs with a single disulfide compared to the dual disulfide stabilized designs (denoted ss2). **B.** TcdB neutralization using2 at 0.1 pM comparing the parental designs (solid) to the dual disulfide stabilized designs (dashed) in the CSPG-dependent system.

**Figure S9.**
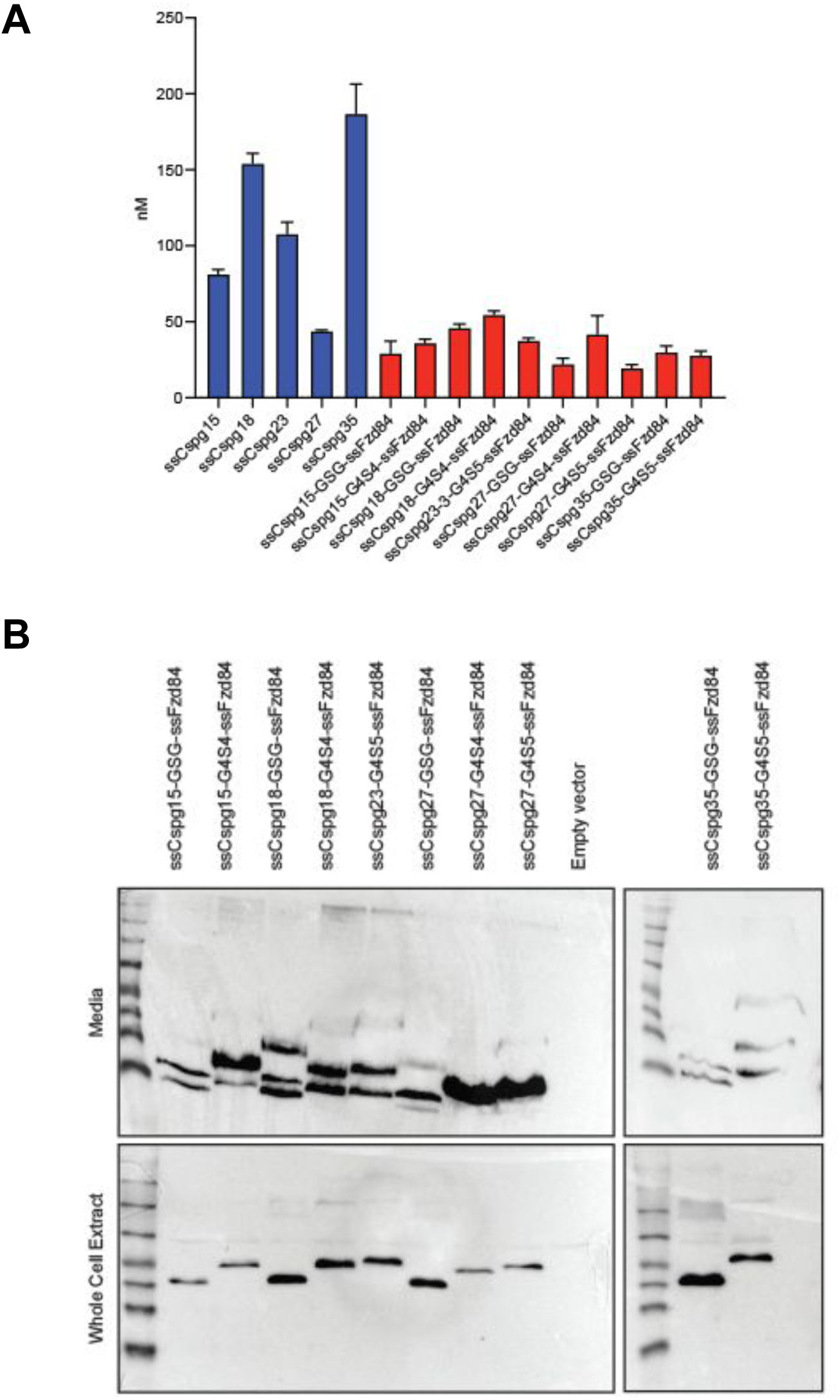
Secretion of fusion binder constructs from *S. boulardii*. **A.** Quantified secretion of monomeric ss2cspg designs and their fused counterparts to ssfzd84 into culture media. B. Western blot showing efficient secretion of the fusion constructs into cell culture media.

## Supplementary Tables

**Table S1:** DIlution series for yeast surface display FACS.

| Target site | Stage | Titration series |
| --- | --- | --- |
| Frizzled | Initial | 1000 nM, 100 nM, 10 nM |
| Frizzled | SSM | 1000 nM, 100 nM, 10 nM, 1 nM |
| Frizzled | Combo | 1 nM, 333 pM, 111 pM, 37 pM |
| CSPG4 | Initial | 1000 nM, 100 nM, 10 nM, 1 nM |
| CSPG4 | SSM | 1000 nM, 100 nM, 10 nM, 1 nM |
| CSPG4 | Combo | 1 nM, 100 pM, 20 pM |

**Table S2:**
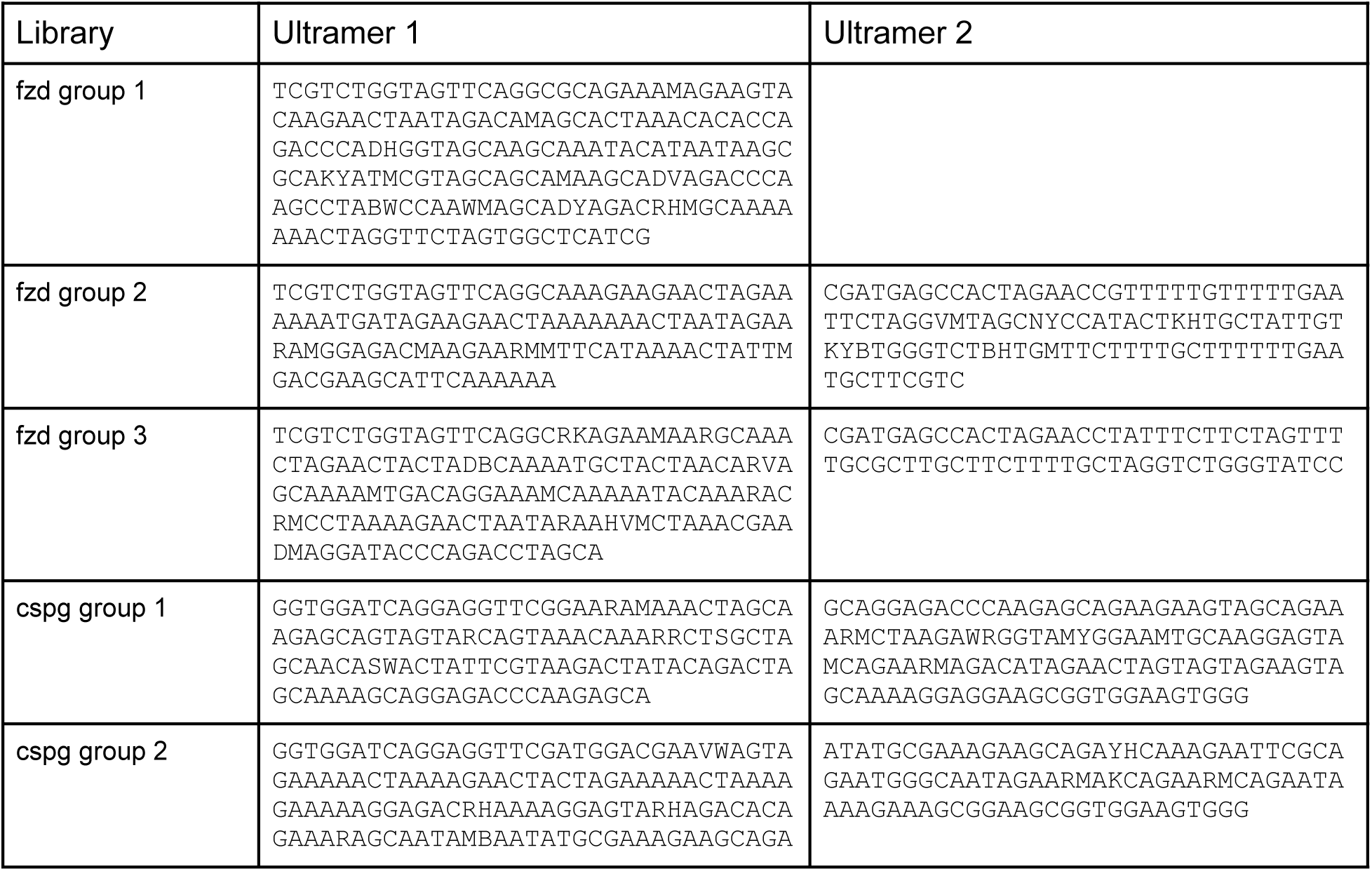
Ultramer sequences for assembly of the combo libraries for fzd and cspg designs.

**Table S3:**
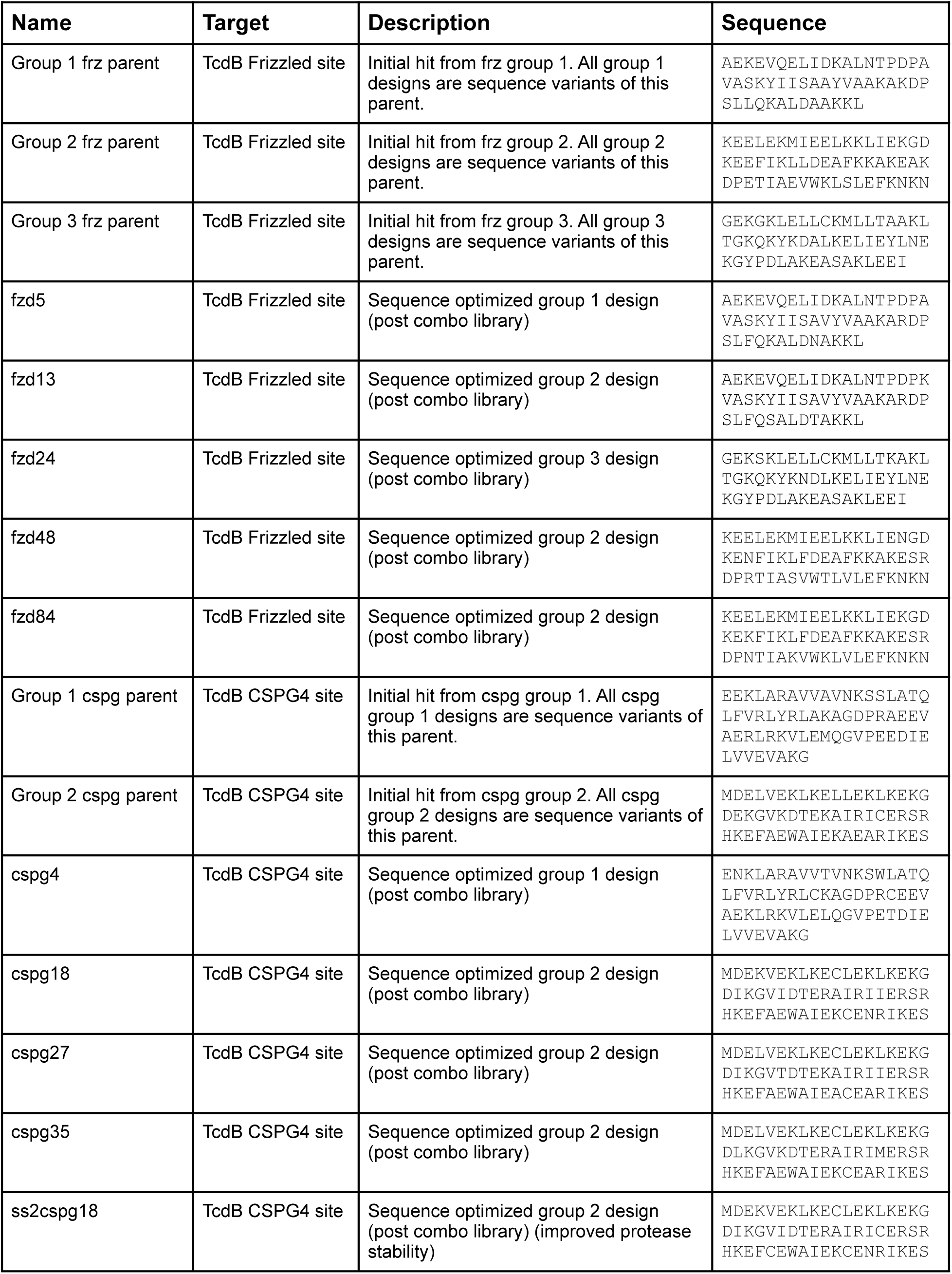
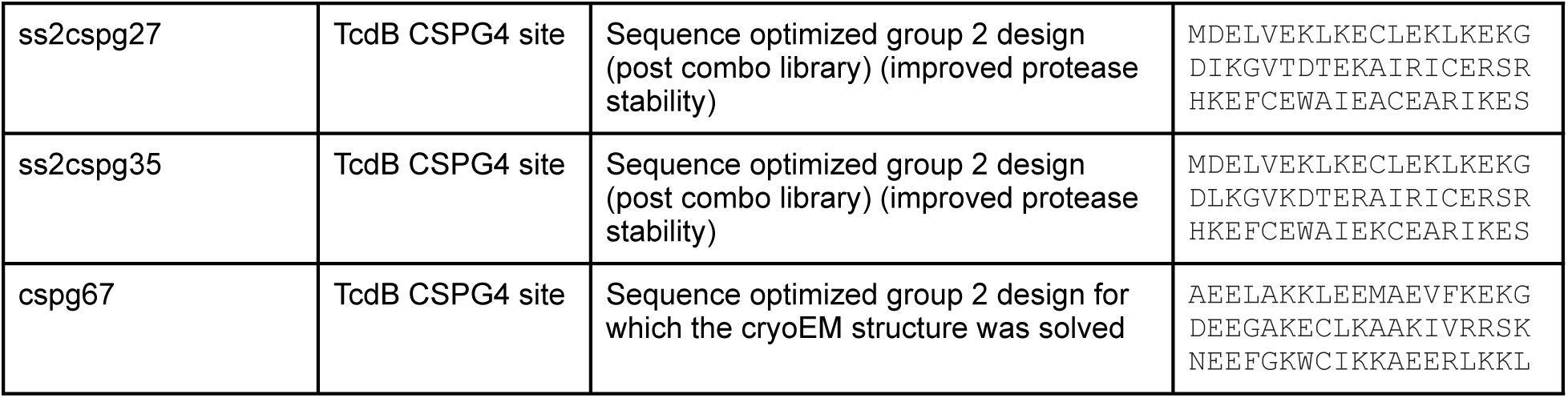
Descriptions and amino acid sequences of key designs described in the paper.

**Table S4:** Cryo-EM data collection, refinement, and validation statistics for TcdB with Frizzled-blocking minibinder.

|  |  |
| --- | --- |
| **CryoEM data collection** |  |
| Magnification | 105,000 |
| Voltage (kV) | 300 |
| Electron exposure | 47 e/Å<sup>2</sup> |
| Pixel size | 2.7 Å |
| Symmetry | C1 |
| Number of particles | 108,076 |
| Map resolution | 4.64 Å |
| FSC threshold | 0.143 |
| **Model Composition – chain A:TcdB** |  |
| Nonhydrogen atoms | 9,354 |
| Protein residues | 1870 |
| Modeled residues of TcdB | 1-1094, 1196-1230,<br>1280-1352, 1373-1432,<br>1453-1986, 1997-2024 |
| Starting model |  |
| **Model Composition – chain B :fzd48** |  |
| Nonhydrogen atoms | 287 |
| Protein residues | 57 |
| **Bonds (RMSD)** |  |
| Bond Length (Å) | 0.010 |
| Bond Angle (°) | 2.165 |
| **Validation** |  |
| MolProbity score | 1.35 |
| Clashscore | 0.14 |
| Poor rotamers | 0 |
| **Ramachandran plot** |  |
| Favored (%) | 75.56 |
| Allowed (%) | 19.31 |
| Disallowed (%) | 5.13 |

**Table S5:**
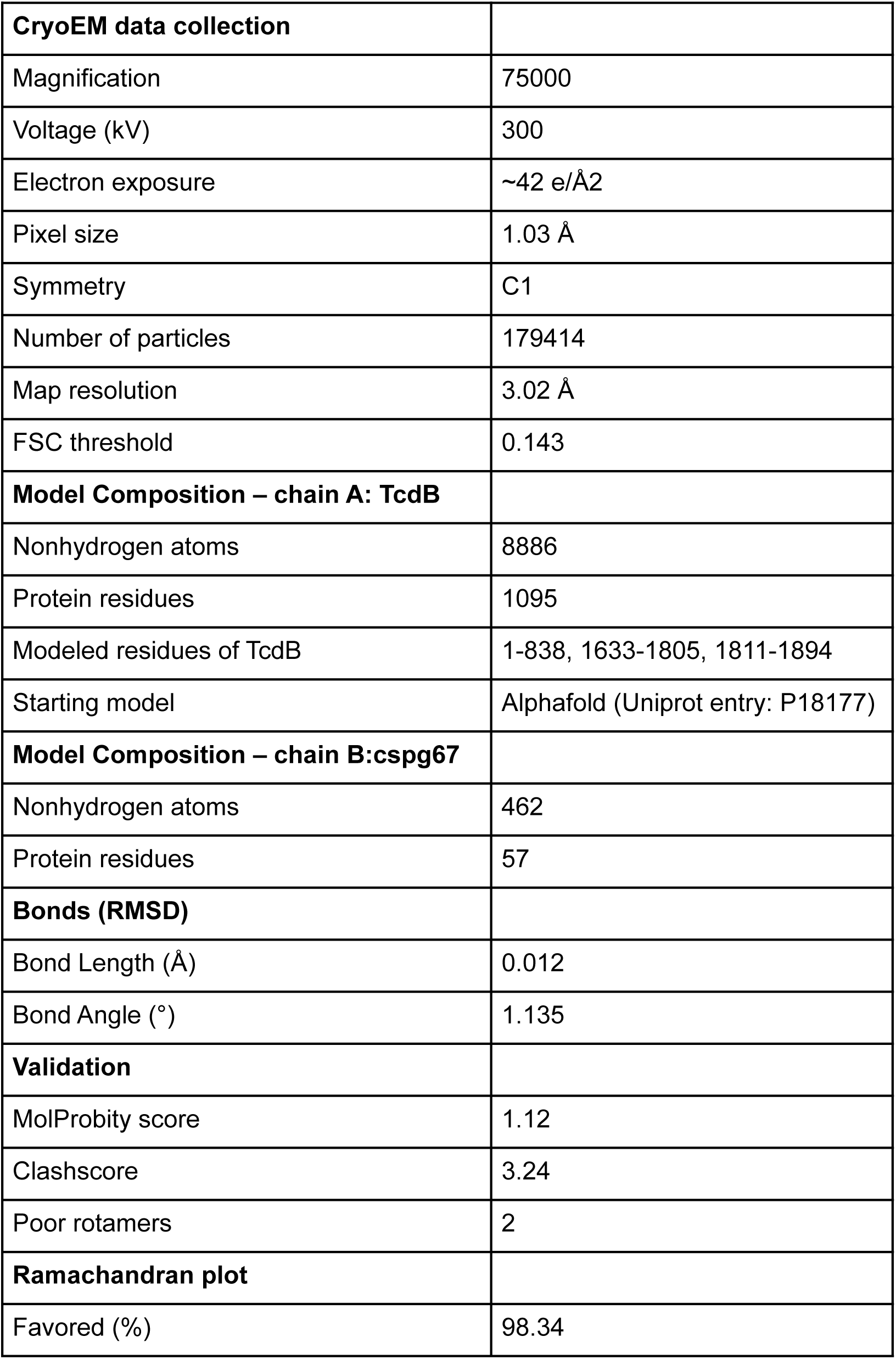
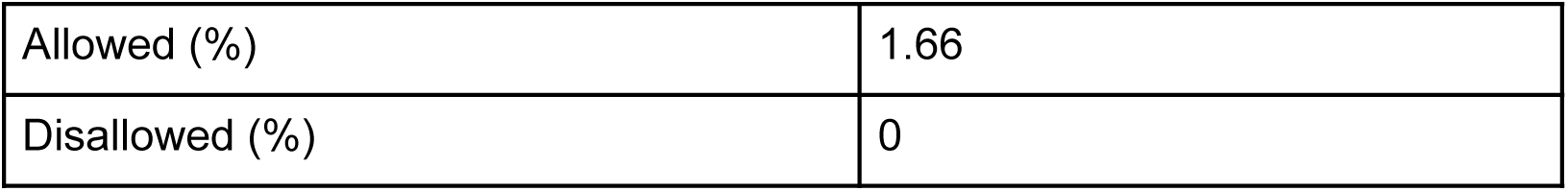
Cryo-EM data collection, refinement, and validation statistics for TcdB with CSPG4- blocking minibinder.

|  |  |
| --- | --- |
| **CryoEM data collection** |  |
| Magnification | 75000 |
| Voltage (kV) | 300 |
| Electron exposure | ~42 e/Å<sup>2</sup> |
| Pixel size | 1.03 Å |
| Symmetry | C1 |
| Number of particles | 179414 |
| Map resolution | 3.02 Å |
| FSC threshold | 0.143 |
| **Model Composition – chain A: TcdB** |  |
| Nonhydrogen atoms | 8886 |
| Protein residues | 1095 |
| Modeled residues of TcdB | 1-838, 1633-1805, 1811-1894 |
| Starting model | AlphaFold (Uniprot entry: P18177) |
| **Model Composition – chain B:cspg67** |  |
| Nonhydrogen atoms | 462 |
| Protein residues | 57 |
| **Bonds (RMSD)** |  |
| Bond Length (Å) | 0.012 |
| Bond Angle (°) | 1.135 |
| **Validation** |  |
| MolProbity score | 1.12 |
| Clashscore | 3.24 |
| Poor rotamers | 2 |
| **Ramachandran plot** |  |
| Favored (%) | 98.34 |
| Allowed (%) | 1.66 |
| Disallowed (%) | 0 |

